# Rewiring the fusion oncoprotein EWS/FLI1 in Ewing sarcoma with bivalent small molecules

**DOI:** 10.1101/2025.03.14.643353

**Authors:** Michael J. Bond, Ryan P. Golden, Giulia DiGiovanni, Briana Howard, Roman C. Sarott, Basel A. Karim, Sai Gourisankar, Gabriela Alexe, Kenneth Ross, Nathanael S. Gray, Kimberly Stegmaier

**Affiliations:** Department of Pediatric Oncology, Dana-Farber Cancer Institute and Boston Children’s Hospital, Boston, MA, USA; The Broad Institute of MIT and Harvard, Cambridge, MA, USA; Department of Chemistry, Stanford University, Stanford, CA, USA; Department of Chemical and Systems Biology, Stanford Cancer Institute, ChEM-H, Stanford University, Stanford, CA, USA

## Abstract

Deregulated transcription is a defining hallmark of cancer, especially pediatric malignancies, which are frequently driven by fusion transcription factors. Targeting transcription factors directly has been challenging as they lack druggable pockets. Recently, chemically induced proximity has enabled the rewiring of transcriptional activators to drive expression of pro-apoptotic genes using bivalent small molecules. Targeting fusion transcription factors, such as EWS/FLI1 in Ewing sarcoma, with these compounds, may open new therapeutic avenues. Here, we develop a small molecule, **EB-TCIP**, that recruits FKBP12^F36V^-tagged EWS/FLI1 to DNA sites bound by the transcriptional regulator BCL6, leading to rapid expression of BCL6 target genes. **EB-TCIP** activity is dependent on ternary complex formation and specific to cells that express FKBP-EWS/FLI1. This proof-of-concept study demonstrates that EWS/FLI1 can be relocalized on chromatin to induce genes that are ordinarily regulated by a transcriptional repressor. Insights herein will guide the development of bivalent molecules that rewire fusion transcription factors.

## Introduction

Over the past two decades, chemical biologists have reshaped how scientists interrogate biological systems by developing tool molecules that hijack numerous enzyme classes^1–7^. Most recently, transcriptional activators have been rewired to drive expression of pro-apoptotic genes using **T**ranscriptional/epigenetic **C**hemical **I**nducers of **P**roximity (TCIPs)^8^. Binders of known transcriptional activators BRD4 and CDK9 were linked to a B cell lymphoma 6 (BCL6) inhibitor to induce expression of pro-death genes leading to apoptosis in a lineage-specific fashion^8,9^. These studies have established TCIPs as a promising new therapeutic modality for cancers whose survival is dependent on suppression of apoptosis^10^. Hijacking fusion transcription factors (TFs) expressed only in tumor cells presents another exciting application of this technology that could be leveraged toward tumor specific therapeutic benefit.

Many cancers, but particularly pediatric malignancies, are driven by fusion TFs that are expressed solely in tumor cells and these cancers have otherwise relatively quiet genomes with few additional genetic abnormalities^11–13^. Therefore, directly targeting the fusion TF could yield potent therapeutic activity with a favorable toxicity profile. Ewing sarcoma (ES) is a solid tumor of the bone that is driven by a single chromosomal translocation, which results in the expression of a fusion TF comprised of the N-terminal transactivation domain of a FUS, EWS, TAF15 (FET) family RNA binding protein fused to the DNA binding domain of an E26 Transformation Specific (ETS) family TF^14^. ETS TFs contain a N-terminal regulatory domain and control expression of genes important for cell growth and survival^15^. In the FET/ETS fusion proteins that arise in ES, the fusion TF retains the ability to bind to canonical ETS target genes but acquires the strong transactivation domain of the FET protein. Moreover, the fusion TF gains the ability to bind long GGAA microsatellite repeats, where it acts as a pioneering TF, opening chromatin and establishing *de novo* enhancers that interact with promoters and boost gene expression^16,17^. The most common FET/ETS fusion results from the (11;22)(q24;q12) translocation, which fuses the Ewing sarcoma breakpoint region 1 (EWSR1) protein to the Friend leukemia integration 1 (FLI1) TF, forming the EWS/FLI1 fusion TF^14^.

EWS/FLI1 accounts for 85% of all ES cases^14^. As a specific and strong dependency in Ewing sarcoma based on CRISPR, RNAi, and degradation-based approaches, EWS/FLI1 should be a prime candidate for drug discovery^13,14^. Unfortunately, the disordered nature of the fusion TF has made it difficult to identify small molecule binders. Due to the dearth of EWS/FLI1-specific ligands, we have used a N-FKBP12^F36V^-EWS/FLI1 (FKBP-E/F) model system to test whether EWS/FLI1 can be relocalized to new sites on chromatin. The FKBP12^F36V^ domain of the FKBP-E/F fusion protein binds specifically and with high affinity to *ortho*-AP1867 (**OAP**)^18^, which can be used as a small molecule handle to hijack FKBP-E/F activity. Given that TCIPs have successfully targeted BCL6 as a transcriptional repressor of interest (i.e., known chemical matter, validated exit vector, and assay availability) and that ES cells express this protein at moderate to high levels (Figure SI-1A), we synthesized and tested a library of bivalent molecules composed of **OAP** linked to **BI3812**, an inhibitor of BCL6. Although BCL6 is well studied in the maturation of B cells and the tumorigenesis of B cell lymphomas, its role in ES biology is less well understood. We first used genomic approaches to identify relevant BCL6 target genes in ES cells. We then used biochemical and omics approaches to characterize the ability of our lead molecule, termed **EB-TCIP**, to induce expression of ES relevant BCL6 targets and compared its activity to the effect of small molecule inhibition and/or degradation of BCL6. Our study demonstrates that EWS/FLI1 can be moved on chromatin to induce expression of neo-target genes, representing the first steps in understanding how the transcriptional machinery of EWS/FLI1 can be reprogrammed for therapeutic effect. Lessons learned from this study may inform future therapeutics for the treatment of TF-fusion driven cancers.

## Results

### Identifying BCL6 Target Genes in ES cells

BCL6 is well-known for its oncogenic role in diffuse large B-cell lymphoma (DLBCL), where it acts as a repressor of *TP53* and associated DNA damage/proapoptotic genes, as well as cell cycle checkpoint genes such as *CDKN1A*^19^. Little is known about the role of BCL6 in ES tumorigenesis, and while BCL6 corepressor (BCOR) fusions are common in Ewing-like sarcomas, not much is known about the role BCL6 plays in these tumors either^20^. Nonetheless, DepMap expression data shows that ES cells express BCL6 at higher levels than in many other cancer types, DLBCL being an exception (Figure SI-1A) ^21^. BCL6 is not a dependency in ES, while it is in DLBCL (Figure SI-1B).

To identify BCL6 target genes in ES cells, we used two *BCL6* targeting guides from the Avana CRISPR guide library^22^ to knockout (KO) *BCL6* in two distinct ES models. Many cultured ES cell lines are *TP53* mutant, even though most patient tumors are *TP53* wild type^13^. Since BCL6 represses *TP53* and related transcripts, we used RNA-seq to profile transcriptional changes in EWS502 (*TP53* mutant) and TC32 (*TP53* wild type) cells. KO of *BCL6* led to few, but consistent changes in RNA transcripts in both models (Figure 1A-B). To verify that the observed signature was related to BCL6, we performed Gene Set Enrichment Analysis (GSEA)^23^ against a gene set derived from *BCL6* promoter binding data generated in primary B cells and DLBCL^24^. We observed a significant, positive correlation between both ES *BCL6* KO models and the published BCL6 repressed gene set (Figure 1C-D). The two transcripts that were the most significantly upregulated in both cell lines were *SOCS2* and *CISH*. These transcripts encode for E3 ligase subunit paralogs involved in the degradation of growth hormone receptor and other cytokine receptors within the JAK/STAT signaling pathway^25,26^. We verified that *SOCS2* and *CISH* transcripts increase with *BCL6* KO by RT-qPCR (Figure 1E-H). Further, SOCS2 protein levels were enhanced upon *BCL6* KO (Figure 1I-J). With *SOCS2* and *CISH* identified as *bona-fide* BCL6 repressed targets, we set out to determine if a TCIP molecule could hijack FKBP-E/F and enhance their expression.

**Figure 1:**
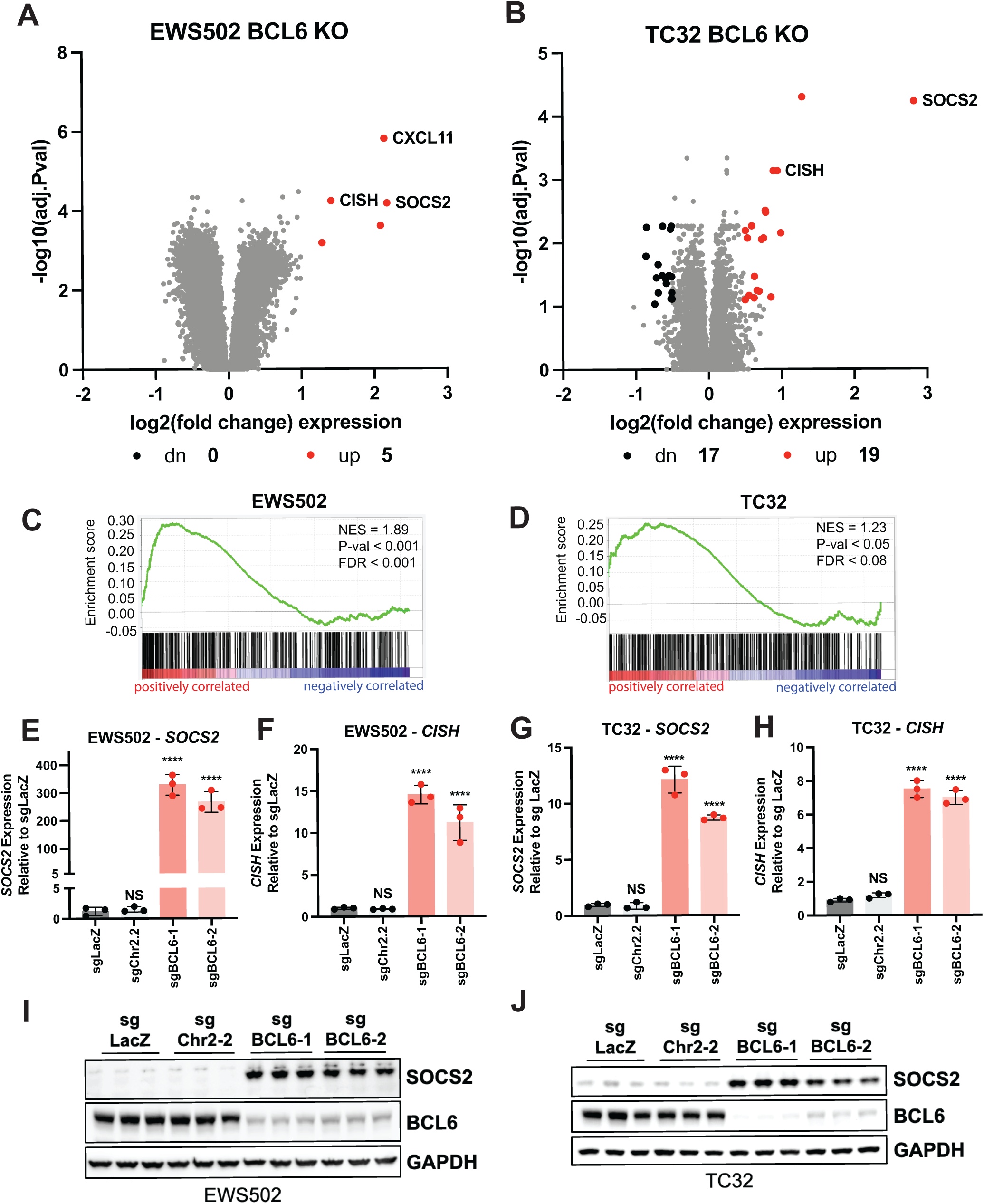
Determining BCL6 target genes in ES. Volcano plots of differentially expressed RNA species in *BCL6* KO (two guides averaged over three replicates per guide) EWS502 cells (A) or TC32 cells (B) vs Chr2.2 control cells. GSEA in EWS502 (C) or TC32 (D) *BCL6* KO cells shows a positive correlation with a published BCL6 gene signature derived from *BCL6* promoter binding data^24^. *SOCS2* and *CISH* transcripts have increased expression by RT-qPCR in *BCL6* KO EWS502 cells (E and F) or TC32 cells (G and H) vs control guides. Average expression from three independent cell transductions (performed in technical triplicate) is shown. RT-qPCR data was compared by one-way ANOVA; NS = not significant, **** p < 0.001. All error bars in the figure show mean ± SD. Immunoblotting shows *BCL6* KO and corresponding increase in SOCS2 protein levels in EWS502 (I) and TC32 (J) cells. Each lane is from an independent transduction of cells. GAPDH serves as a loading control.

### EB-TCIP induces BCL6 target gene expression more effectively than chemical inhibition or degradation of BCL6

Although EWS/FLI1 has been a priority target for ES drug discovery, there has been limited success identifying EWS/FLI1 ligands. EWS/FLI1, like many TFs, is highly disordered and difficult to drug. To overcome this issue, we took advantage of N-terminal tagged FKBP12^F36V^-EWS/FLI1 (FKBP-E/F) cell lines that were previously developed to study EWS/FLI1 degradation using the dTAG system^27,28^. We envisioned that **BAK-04-212**, a bivalent molecule comprised of **OAP** and **BI3812**, which we call **EB-TCIP**, could redirect FKBP-E/F to BCL6 loci, thereby driving expression of BCL6 repressed transcripts such as *SOCS2* and *CISH* (Figure 2A-B).

**Figure 2:**
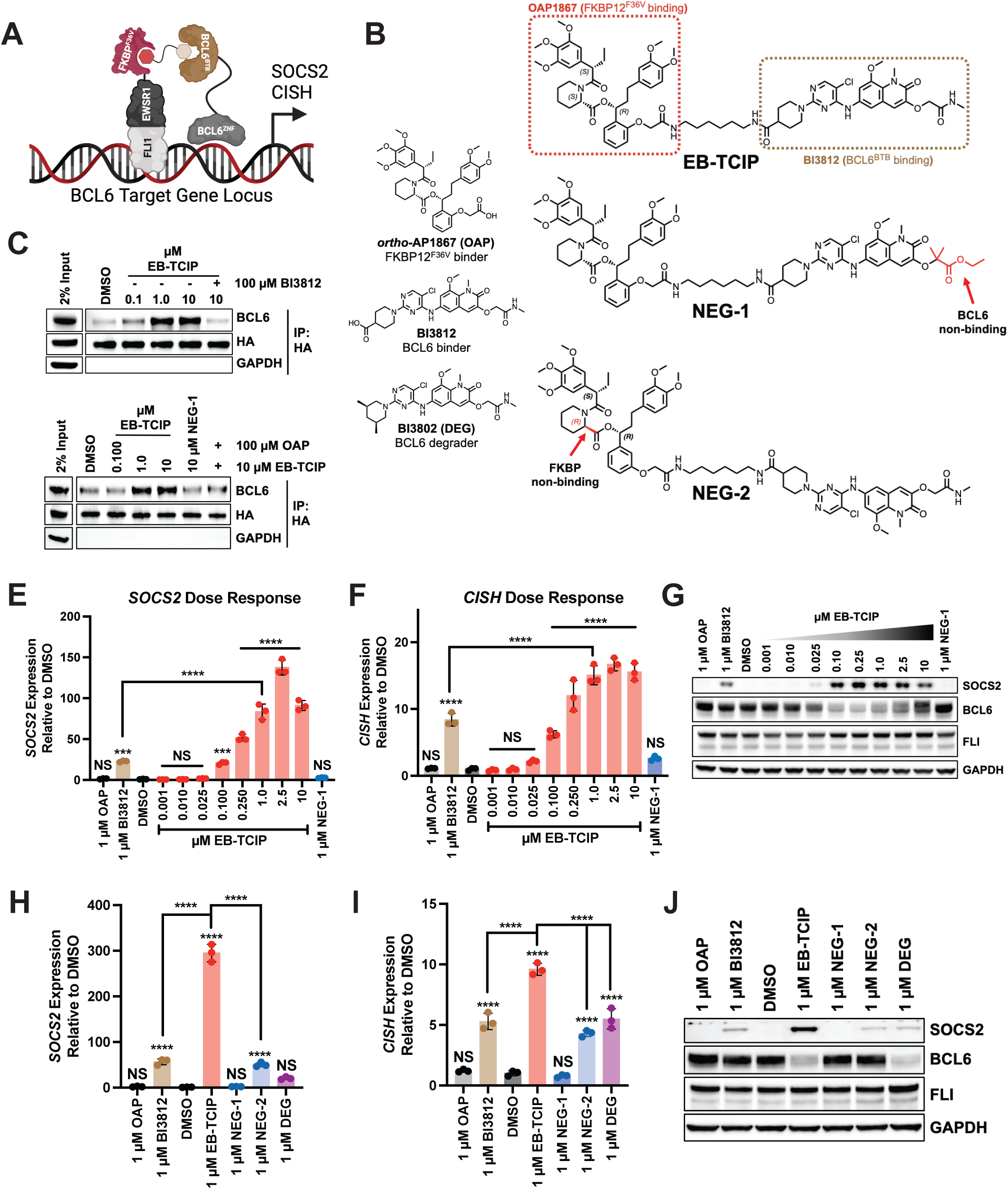
EB-TCIP increases expression of BCL6 target genes in ES cells at nanomolar concentrations. (A) Schematic of **EB-TCIP** mechanism of action. **EB-TCIP** induces a ternary complex between FKBP^F36V^ tagged EWS/FLI1 and BCL6, which leads to activation of BCL6 target gene transcription. Image made with Biorender. (B) Structures of compounds used in this work. (C) **EB-TCIP** increases the association of BCL6 with **EB-TCIP** in a dose dependent manner in EWS502 FKBP-E/F cell lysates while **NEG-1** (D) does not induce a ternary complex. The association is reversible as excess **BI3812** (C) and excess **OAP** (free acid) (D) abrogate ternary complex formation. GAPDH was probed to determine if unbound proteins were removed by washing. **EB-TCIP** dose dependently increases *SOCS2* (E) and *CISH* (F) expression by RT-qPCR. (G) SOCS2 protein levels dose dependently increase while BCL6 protein levels dose dependently decrease after **EB-TCIP** treatment. **EB-TCIP** induces higher *SOCS2* (H) and *CISH* (I) transcript levels than chemical inhibition with **BI3812** or chemically induced degradation with **BI3802 (DEG)**. (J) SOCS2 protein levels are highest in **EB-TCIP** treated cells compared to **BI3812**, **BI3802 (DEG)**, or negative control compounds that do not form ternary complexes. Immunoblotting is representative of three biological replicates and GAPDH serves as a loading control. All experiments were performed in FKBP-E/F expressing EWS502 cells. RT-qPCR experiments show one experiment with technical triplicate that is representative of three biological replicates. Means of *SOCS2* and *CISH* expression were compared using one-way ANOVA with multiple comparisons; NS = not significant, *** p < 0.005, **** p < 0.001. All error bars in the figure indicate mean ± SD. Unless indicated with brackets, significance above each condition indicates comparison of that mean to the mean of DMSO.

We first wanted to demonstrate that **EB-TCIP** can induce a ternary complex between FKBP-E/F and BCL6 in a cell free system using time-resolved fluorescence energy transfer (TR-FRET) between the BTB domain of BCL6 (BCL6^BTB^) labelled with fluorescein isothiocyanate (FITC) and a His-tagged FKBP^F36V^, which was recognized by an anti-His-tag terbium-conjugated antibody^29^. **EB-TCIP** dose dependently increased TR-FRET signal with an EC_50_ of 0.14 ± 0.03 µM, while the negative control bifunctional compound **RPG-02-089**, referred to as **NEG-1**, did not increase TR-FRET signal (Figure SI-2A). The addition of two vicinal methyl groups in **NEG-1** sterically occludes binding to the BCL6^BTB^. At concentrations of **EB-TCIP** above 0.31 µM a hook effect was observed. This is a characteristic property of bivalent molecules where at high concentrations, binary complexes between the compound and one target predominate over the ternary complex^30^. Next, we investigated formation of a ternary complex between FKBP-E/F and native BCL6. To this end, we treated EWS502 FKBP-E/F cell lysates with increasing concentrations of **EB-TCIP**. Since the FKBP-E/F construct contains a HA tag, we then used magnetic HA beads to immunoprecipitate FKBP-E/F and associated proteins. We observed a dose dependent increase in the amount of BCL6 pulled down in the presence of **EB-TCIP** (Figure 2C-D). Further, **NEG-1** was unable to pulldown BCL6. By pre-treating lysates with either excess **BI3812** or **OAP** before **EB-TCIP** treatment, the ternary complex was disrupted and little BCL6 was pulled down compared to treatment with 1 or 10 µM **EB-TCIP** alone (Figure 2C-D). These data demonstrate that **EB-TCIP** can form a reversible ternary complex between FKBP^F36V^ and BCL6^BTB^ *in vitro* and in cell lysates.

After confirmation of ternary complex formation, we next tested if **EB-TCIP** could enhance expression of BCL6 repressed targets. Previous TCIP studies monitored compound activity using a BCL6 repressed GFP reporter (Figure SI-2B)^8,9^. Using this same vector, we engineered an EWS502 FKBP-E/F line expressing the reporter and found that **EB-TCIP** dose dependently increased the percentage of GFP positive cells to a greater extent than negative control compounds (Figure SI-2C). Next, we treated EWS502 FKBP-E/F cells with increasing concentrations of **EB-TCIP** and monitored expression of identified BCL6 targets by RT-qPCR and immunoblotting. **EB-TCIP** dose dependently increased expression of *SOCS2* and *CISH*, with an EC_50_ of 0.17 ± 0.05 µM and 0.11 ± 0.04 µM respectively (Figure 2E-F). Maximal induction of both transcripts was reached at a concentration of 2.5 µM with a hook effect evident at 10 µM. **EB-TCIP** induced higher levels of expression of both transcripts compared to **BI3812** at 1 µM. Additionally, 1 µM of **NEG-1** did not increase *SOCS2* or *CISH* expression. A dose dependent increase in SOCS2 protein level was also observed (Figure 2G). **EB-TCIP** induced higher SOCS2 protein expression than **BI3812** and **NEG-1**. Similar trends in transcript and protein expression were observed for TC32 FKBP-E/F cells (Fig S-2D-F, Table S1) demonstrating the activity of **EB-TCIP** is not unique to EWS502 FKBP-E/F cells. **EB-TCIP** dose dependently decreased proliferation of EWS502 FKBP-E/F cells over 72 hours (Figure SI-2G). However, we also observed similar antiproliferative activity in parental EWS502 cells that do not express exogenous FKBP-E/F (Figure SI-2H). Our viability data suggests **EB-TCIP** induces off-mechanism cytotoxicity. Nonetheless, at shorter timepoints **EB-TCIP** is a useful tool molecule to study relocalization of FKBP-E/F on chromatin.

We observed a dose dependent decrease in BCL6 protein levels in both EWS502 and TC32 FKBP-E/F cells at concentrations where SOCS2 levels increase and ternary complex between FKBP-E/F and BCL6 is formed. The decrease in BCL6 protein was not due to a decrease in BCL6 transcript levels as treatment with **EB-TCIP** increased BCL6 mRNA (Figure SI-2I), which is consistent with previous TCIP studies^8^. We wondered if BCL6 induced degradation was enough to increase SOCS2 and CISH protein/transcript levels to the same extent as **EB-TCIP**. Therefore, we treated cells with **BI3802**^31^ (Figure 2B), which induces the polymerization and subsequent proteasome dependent degradation of BCL6^32^. **BI3802** induced BCL6 degradation to a similar extent as **EB-TCIP**; however, **EB-TCIP** induced significantly higher levels of *SOCS2* and *CISH* transcripts, as well as SOCS2 protein, compared to **BI3802** (Figure 2H-J).

During our characterization of **EB-TCIP**, the synthesis of an **OAP** derivative that does not bind to FKBP12^F36V^ was described^33^. Using this synthesis, we generated a second negative control compound, **RPG-02-205**, referred to as **NEG-2** (Figure 2B), that does not form a ternary complex but retains the ability to engage BCL6. **NEG-2** did not increase BCL6 target gene/protein expression to the same extent as **EB-TCIP** (Figure 2H-J). The activity that was observed can be attributed to **NEG-2**’s retained ability to inhibit BCL6. **NEG-2** also did not induce BCL6 degradation, providing evidence that **EB-TCIP** decreases BCL6 protein levels in a FKBP-E/F dependent manner.

Our observations above led us to hypothesize that **EB-TCIP** induces proteasome dependent degradation of BCL6. We tested this hypothesis by pre-treating EWS502 FKBP-E/F cells for 1 h with the proteasome inhibitor MG132^34^ or the neddylation inhibitor MLN4924^35^ before treatment with **BI3802** or **EB-TCIP** for 4 h. MG132 rescued BCL6 levels to a greater extent than MLN4924, which was seen previously for **BI3802**^32^ (Figure SI-3A). As a control we also pre-treated cells with the transcriptional inhibitor Actinomycin D^36^ (ActD). ActD treatment abrogated **EB-TCIP** activity as expected (Figure SI-3A-C). These data suggest **EB-TCIP** activity is dependent on both active transcriptional and degradation machinery. We propose a mechanism by which a protein associated with FKBP-E/F induces BCL6 degradation, allowing FKBP-E/F to bind chromatin and drive transcription of BCL6 targets (Figure SI-3D).

### EB-TCIP induces rapid, ternary complex dependent induction of BCL6 targets that is specific to cells expressing FKBP-E/F

To determine the kinetics of BCL6 degradation and target induction we treated EWS502 FKBP-E/F cells with DMSO, **BI3812**, **EB-TCIP**, or **BI3802** (1 µM) over a 24 h time course. **EB-TCIP** and **BI3802** induced rapid degradation of BCL6 with near maximal degradation observed within 1 h (Figure 3A). Despite similar degradation kinetics, **EB-TCIP** enhanced SOCS2 protein levels to a greater extent than **BI3802** at all time points beyond 2 h. **EB-TCIP** also enhanced SOCS2 protein levels more than **BI3812** at these time points. Protein expression lagged behind transcript expression, which at 1 h was significantly higher in **EB-TCIP** treated cells compared to DMSO or the other molecules (Figure 3B-C). **EB-TCIP**-induced expression of *SOCS2* and *CISH* showed a peak between 2 and 4 h. *SOCS2* expression levelled off before increasing at 24 h, whereas *CISH* expression continued to decrease until the end of the experiment. Increases in these transcripts in **BI3812** and **BI3802** treated cells were relatively stable after 2 h, suggesting **EB-TCIP** has a different mechanism of transcript induction than these compounds. These data show that **EB-TCIP** rapidly and more effectively induces expression of BCL6 targets compared to chemical inhibition or degradation.

**Figure 3:**
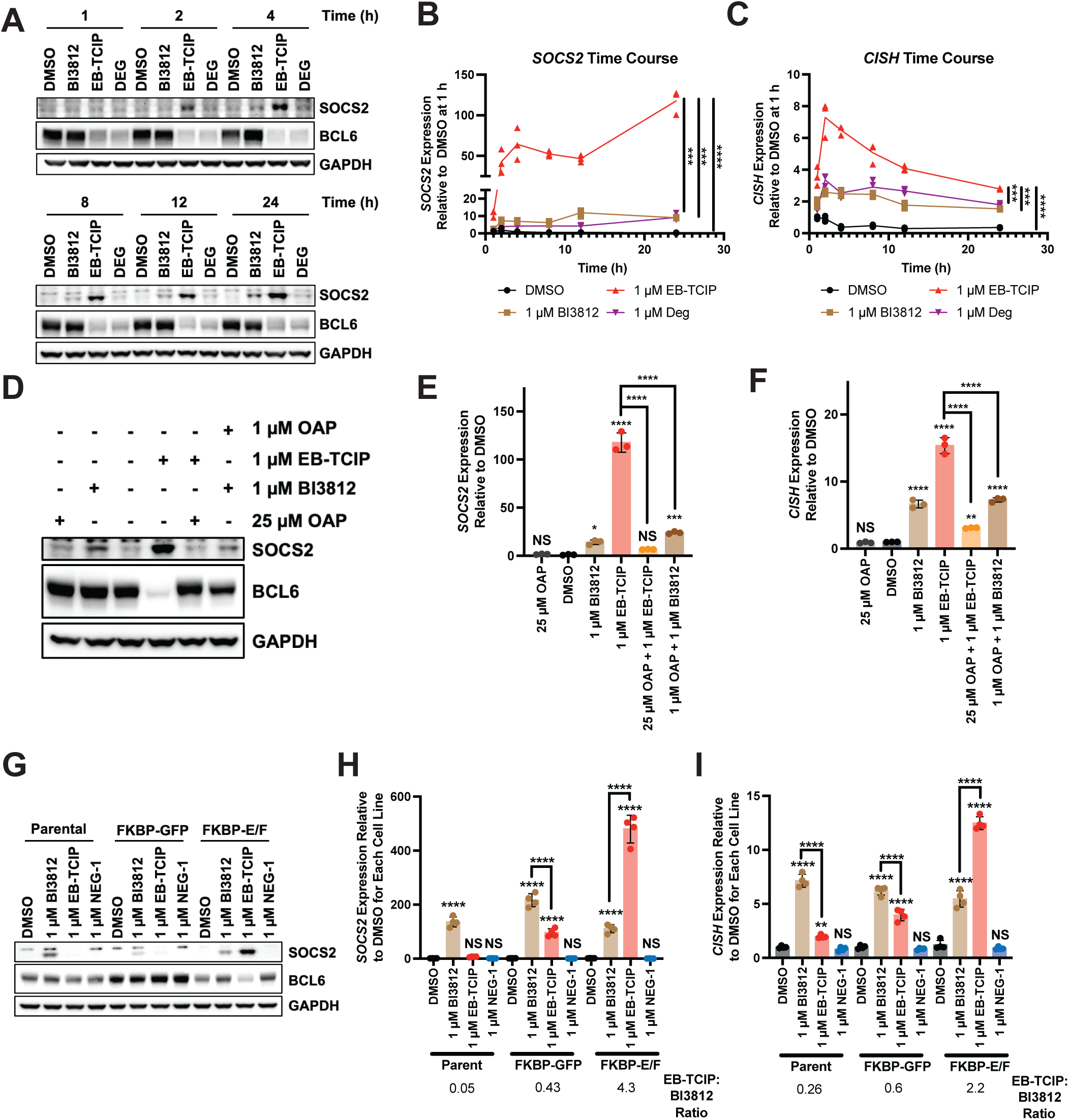
EB-TCIP activity is rapid, ternary complex dependent, and specific to cells expressing FKBP-E/F. (A) Time course of SOCS2 and BCL6 protein levels. BCL6 degradation occurs within 1 h for both **EB-TCIP** and **BI3802 (DEG)**. **EB-TCIP** induces SOCS2 expression by 2 h and maintains higher expression levels than **BI3812** or **BI3802 (DEG)** throughout the time course. *SOCS2* (B) and *CISH* (C) transcripts reach a maximum between 2 and 4 h by RT-qPCR. (D) **EB-TCIP** induced SOCS2 protein expression can be reversed with 25-fold excess **OAP** (free acid). Co-treatment of 1 µM **BI3812** and **OAP** do not increase SOCS2 protein expression more than 1 µM **BI3812** alone. **EB-TCIP** induced *SOCS2* (E) and *CISH* (F) transcript expression is reversed with excess **OAP**. **BI3812** and **OAP** must be chemically linked to induce maximum transcript expression. (G) **EB-TCIP** induces the highest expression of SOCS2 protein in EWS502 FKBP-E/F cells compared to EWS502 parental cells or EWS502 cells expressing FKBP-GFP. Only treatment with **EB-TCIP** induces more expression of *SOCS2* (H) and *CISH* (I) than **BI3812** in EWS502 FKBP-E/F cells. **EB-TCIP**:**BI3812** ratio was calculated by dividing the average expression of each transcript in **EB-TCIP** treated cells by the average expression of each transcript in **BI3812** treated cells. Immunoblotting is representative of three biological replicates. RT-qPCR experiments show one experiment with technical triplicate or quadruplicate that is representative of three biological replicates. The means of *SOCS2* and *CISH* expression were compared using one-way ANOVA with multiple comparisons; NS = not significant, * p < 0.05, ** p < 0.01, *** p < 0.005, **** p < 0.001. All error bars in the figure represent mean ± SD. Unless indicated with brackets, significance above each condition indicates comparison of that mean to the mean of DMSO. In (H) and (I), unless indicated with brackets, the means of **BI3812**, **EB-TCIP**, and **NEG-1** were compared to the DMSO sample for the corresponding cell line.

Next, we wanted to ensure the activity of **EB-TCIP** was via a ternary complex mechanism. To do this, we pre-treated EWS502 FKBP-E/F cells with a 25-fold excess of **OAP** for 1 h before treating cells with **EB-TCIP** for 4 h. The excess **OAP** competed away **EB-TCIP** and abolished its ability to induce BCL6 target expression (Figure 3D-F). Further, co-treatment with 1 µM of **BI3812** and **OAP** did not increase BCL6 target expression as much as **EB-TCIP** (Figure 3D-F). These data show that **BI3812** and **OAP** must be chemically linked to induce a ternary complex and drive BCL6 target gene expression. To further validate the importance of ternary complex formation and show compound specificity, we tested the ability of **EB-TCIP** to increase BCL6 targets in parental, FKBP-GFP, and FKBP-E/F expressing EWS502 lines. **BI3812** induced SOCS2 protein expression in all cell lines as expected; however, **EB-TCIP** induced SOCS2 protein expression only in FKBP-E/F expressing cells (Figure 3G). Further, BCL6 target transcript levels were highest in FKBP-E/F expressing cells treated with **EB-TCIP** (Figure 3H-I). As a measure of **EB-TCIP** activity we compared the ratio of induction of transcript expression in samples treated with **EB-TCIP** or **BI3812** for each cell line. We observed a positive ratio, indicative of higher activity of **EB-TCIP** than **BI3812**, only in FKBP-E/F expressing cells. Together, these data show the enhanced ability of **EB-TCIP** to induce BCL6 target expression is dependent on ternary complex formation and FKBP-E/F expression.

### EB-TCIP induces rapid, dynamic changes in global transcription

To profile how ternary complex formation between FKBP-E/F and BCL6 affected transcription in an unbiased manner, we treated EWS502 FKBP-E/F cells with DMSO, **BI3812**, **EB-TCIP**, or **NEG-1** (2.5 µM) for 8 or 24 h and studied transcriptomic changes by RNA-seq. Given EWS/FLI1’s ability to activate transcription, at 8 h we observed many more upregulated genes (71) than downregulated genes (4) (Figure 4A). Highly upregulated genes that we observed after *BCL6* KO, such as *SOCS2, CISH, and CXCL11*, were significantly upregulated at 8 h. The number of both upregulated (244) and down regulated genes (116) increased at 24 h with many increasing in magnitude (Figure 4B). However, some genes that were significantly upregulated at 8 h, such as *CISH*, had decreased expression at 24 h, consistent with our earlier time course data.

**Figure 4:**
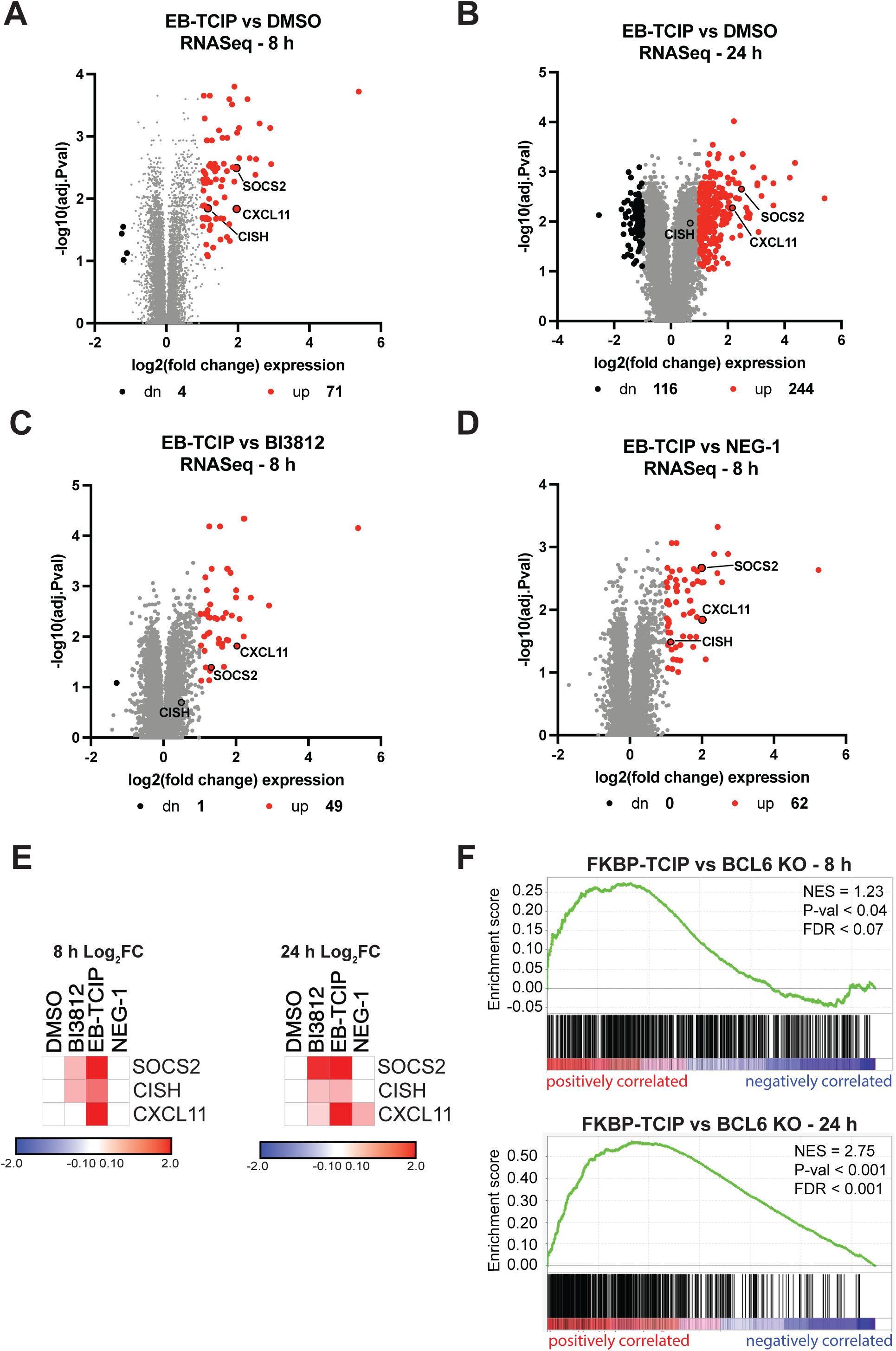
Global RNA changes induced by EB-TCIP are similar to genetic KO of *BCL6* in EWS502 cells. Volcano plots portraying log_2_fold changes of gene expression from cells treated with 2.5 µM **EB-TCIP** versus DMSO at 8 (A) and 24 (B) h with a -log_10_ adjusted P-value cut off of 1. **EB-TCIP** treatment predominantly increases expression of transcripts at both timepoints. Volcano plots portraying log_2_ fold changes of cells treated with 2.5 µM **EB-TCIP** versus 2.5 µM **BI3812** (C) or 2.5 µM **NEG-1** (D) at 8 hours with a -log_10_ P-value cut off of 1. **EB-TCIP** induces higher expression of BCL6 transcripts than **BI3812** or **NEG-1** at this early timepoint. Dots corresponding to *SOCS2, CISH, and CXCL11* are labelled with black borders. (E) Heatmaps of changes in BCL6 target gene expression at 8 (left) and 24 h (right) show that **EB-TCIP** induces faster and/or higher expression of these select genes. (F) GSEA comparing EB-TCIP treated EWS502 FKBP-E/F cells to *BCL6* KO EWS502 parental cells at 8 (top) and 24 h (bottom) show significant positive correlation between the two gene sets. RNA-seq data is shown as the average of three independent replicates.

Global transcriptomic changes were more robust with **EB-TCIP** in comparison to DMSO, **BI3812**, and **NEG-1** (Figure 4C-D and Figure SI-4A-F). **EB-TCIP** induced the expression of more genes than both **BI3812** and **NEG-1** at 8 h. The known BCL6 targets *SOCS2* and *CXCL11* were more upregulated by **EB-TCIP** than **BI3812** or **NEG-1** at 8 h. *CISH* was significantly upregulated by **EB-TCIP** compared to **NEG-1**, but not **BI3812** at 8 h although expression did trend upwards (Figure 4C-E). Conducting RNA-seq at an earlier timepoint may capture the kinetic difference in *CISH* expression between **EB-TCIP** and **BI3812** that we observed in our previous time course. To further asses BCL6 programming induced by **EB-TCIP** we performed GSEA comparing up-regulated genes induced by **EB-TCIP** and the transcriptional changes that result from *BCL6* KO. At both 8 and 24 h, we observed a significant positive correlation between **EB-TCIP** induced gene expression and *BCL6* KO, with *SOCS2*, *CISH*, and *CXCL11* being leading-edge genes within the enriched signature (Figure 4F). Further, at both 8 and 24 h, we observed a significant positive correlation between **EB-TCIP** induced gene expression and the previously published BCL6 target gene set^24^ (Figure SI-4G-H). These data further support that **EB-TCIP** rapidly enhances BCL6 target genes compared to chemical inhibition.

### EB-TCIP relocalizes FKBP-E/F to BCL6 sites on chromatin

To better understand the gene expression changes induced by **EB-TCIP**, we used chromatin immunoprecipitation with sequencing (ChIP-seq) to determine how **EB-TCIP** treatment changes FKBP-E/F and BCL6 localization on chromatin. EWS502 FKBP-E/F cells were treated with DMSO, **BI3812**, **BI3802** or **EB-TCIP** (1 µM) for 24 hours and then subjected to ChIP-seq, using antibodies for HA or BCL6. A HA antibody was used instead of a FLI1 antibody to ensure only FKBP-E/F, and not endogenous EWS/FLI1, was immunoprecipitated. Globally, treatment with all compounds modestly increased FKBP-E/F on chromatin to varying degrees (Figure SI-5A). As expected, degradation of BCL6 induced by **BI3802** and **EB-TCIP** decreased BCL6 binding globally to chromatin, while inhibition with **BI3812** had minimal effect (Figure SI-5B). We observed ∼50% overlap between FKBP-E/F and BCL6 binding sites in DMSO treated HA and BCL6 samples (Figure SI-5C-E), and accordingly, similarities in the binding motifs of EWS/FLI1 and BCL6 (Figure SI-5G). EWS/FLI1 binds DNA at “GGAA” repeats, and this sequence is present within the recognition motif of BCL6. Further, the sequence similarity may enable FKBP-E/F to bind BCL6 target genes with greater affinity when brought into proximity by **EB-TCIP**.

To determine chromatin changes specific to **EB-TCIP** treatment, we clustered peaks in all treatments based on decreasing, unchanged, and increasing peak intensity between **EB-TCIP** and DMSO for both antibodies. This clustered analysis revealed that **EB-TCIP** induced an increase in a subset of both HA and BCL6 peaks, to a greater extent than that observed in **BI3812** or **BI3802** treated samples (Figure 5A and Figure SI-6A). We performed motif analysis to determine what DNA sequences were associated with the **EB-TCIP** treated HA increased peaks and compared this to motif analysis from global HA binding peaks in DMSO treated samples. In DMSO treated samples, the top two motifs were EWS/FLI1 related, as expected (Figure 5B) and no BCL6 motif was observed. However, the BCL6 motif was enriched in HA increased peaks and ranked 29^th^ (Figure 5C). Since **EB-TCIP** induces a ternary complex between FKBP-E/F and BCL6, we explored if the BCL6 increased peaks enriched for EWS/FLI1 signatures. BCL6 increased peaks showed enrichment in EWS/FLI1 motifs and a decrease in the rank of the BCL6 motif compared to DMSO peaks (Figure SI-6B-C).

**Figure 5:**
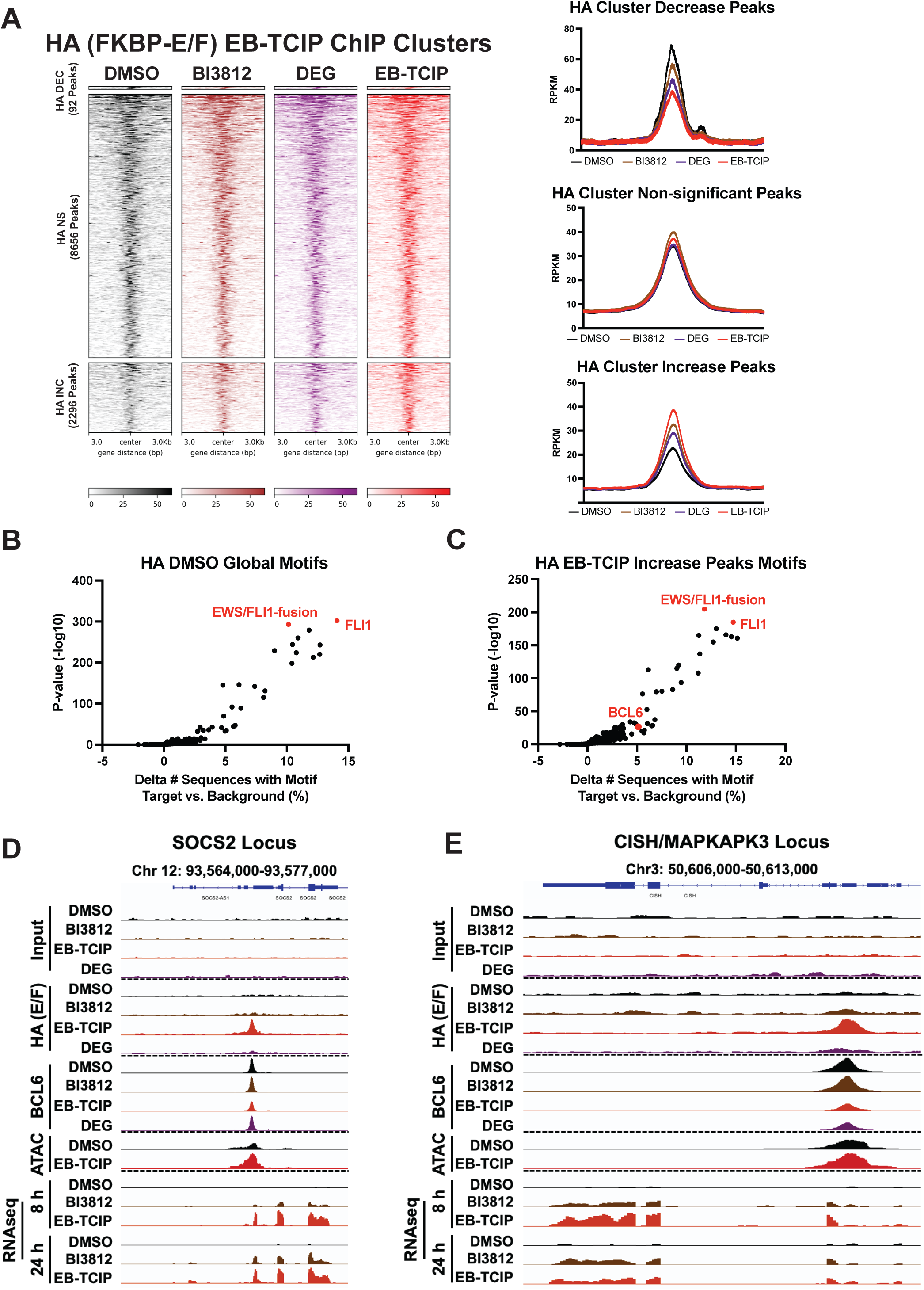
EB-TCIP changes the localization of FKBP-EWS/FLI1 on chromatin. (A) ChIP-seq tornado plots of HA (FKBP-E/F) binding signal of **EB-TCIP** (red) versus DMSO (black) peaks that are decreasing (DEC; 92), non-significantly changing (NS; 8656), and increasing (INC; 2296). Differential peaks between **EB-TCIP** and DMSO are shown for all compounds. Compared to **BI3812** (brown) and **BI3802** (**DEG**, purple), **EB-TCIP** increases FKBP-E/F binding at a subset of genes. Line plots for all compound treatments in each cluster are shown to the right. (B) Scatter plot portraying top enriched motifs of HA binding sites in DMSO treated cells. (C) Scatter plot portraying top motifs of HA increased peaks enriched in **EB-TCIP** treated cells. The BCL6 motif scores 29^th^. IGV visualization of Input, HA (FKBP-E/F), BCL6, ATAC-seq signal, and RNA-seq signal at the *SOCS2* (D) and *CISH* (E) with treatments DMSO (black), 1 µM **BI3812** (brown), 1 µM **BI3802 (DEG**, purple), and 1 µM **EB-TCIP** (red). All ChIP-seq and ATAC-seq is portrayed as the average of two independent replicates. RNAseq is portrayed as the average of three independent replicates.

To understand how changes in DNA binding may impact transcription globally, we compared log_2_fold expression of genes from our 8 h RNA-seq experiment where FKBP-E/F or BCL6 binding changed in the ChIP-seq experiments. Genes from HA increased peaks displayed increased gene expression more so at genes where both FKBP-E/F and BCL6 were bound compared to genes where only FKBP-E/F was bound. Gene expression was similar at HA decreased and HA unchanged peaks regardless of whether BCL6 was bound (Figure SI-6D). Genes from BCL6 increased peaks did not show a decrease in expression, suggesting that FKBP-E/F transcriptional activation was stronger than BCL6 gene repression (Figure SI-6E).

We next visualized changes in FKBP-E/F and BCL6 binding at both BCL6 and EWS/FLI1 target gene sites. Robust BCL6 peaks were observed at *SOCS2, CISH*, and *CXCL11* loci in all treatments (Figure 5D-E and Figure SI-7A). FKBP-E/F binding was not observed in DMSO treated cells and only **EB-TCIP** was able to induce binding. Degradation alone does not explain increased FKBP-E/F binding as **BI3802** and **EB-TCIP** induce similar levels of BCL6 loss at *CISH*, but FKBP-E/F binding is only induced by **EB-TCIP**. Moreover, **EB-TCIP** induced FKBP-E/F binding was specific to BCL6 target loci as no FKBP-E/F was observed at the *GAPDH* genomic locus (Figure-S7-B). For visualization of changes at EWS/FLI1 target sites we focused on *NR0B1 and VRK1*, where EWS/FLI1 canonically binds at a proximal and distal enhancer respectively (Figure SI-8A-B). We observed strong FKBP-E/F binding at both enhancer sites with all treatments. However, only **EB-TCIP** treated samples showed increases in BCL6 binding, which mirrored the distinct pattern of FKBP-E/F at each site. Together, our ChIP-seq data shows that **EB-TCIP**, but not **BI3812** or **BI3802**, can relocalize both FKBP-E/F and BCL6 on chromatin.

Recently, small molecules have been used to redirect the pioneering TF activity of FOXA1 on chromatin^37^. Since **EB-TCIP** relocalizes FKBP-E/F, which has pioneering TF activity, we used assay for transposase-accessible chromatin with sequencing (ATAC-seq) to determine if chromatin accessibility is changed at genomic loci where FKBP-E/F is gained. Globally, chromatin accessibility is not significantly changed with **EB-TCIP** treatment (Figure SI-8C). However, at BCL6 target sites where FKBP-E/F is gained, such as *SOCS2, CISH, and CXCL11*, open chromatin is increased leading to increased RNA-seq peaks (Figure 5D-E and SI-7A). We also investigated changes in chromatin accessibility at BCL6 gained sites *NR0B1* and *VRK1*. EWS/FLI1 binding is relatively unaffected at these sites, as is chromatin accessibility and gene expression (Figure SI-8-A-B). Our ATAC-seq data suggests that relocalized FKBP-E/F increases gene expression by opening chromatin while relocalization of BCL6 is generally not sufficient to repress EWS/FLI1 target genes.

## Discussion

Although ES is the second most common bone cancer in children and adolescents, therapeutic development has been stagnant for decades. In our proof-of-concept study, we demonstrate that the pioneering TF activity of EWS/FLI1 can be redirected to genes typically inactivated by the repressor BCL6. Our tool compound **EB-TCIP**, which links **OAP** to **BI3812**, relocalized FKBP-E/F to chromatin sites bound by BCL6, thereby driving expression of genes ordinarily repressed by BCL6. The compound is potent and induces rapid transcript and protein expression of *SOCS2* and *CISH*. ChIP-seq and ATAC-seq showed that **EB-TCIP** increases open chromatin at FKBP-E/F gained sites. We foresee **EB-TCIP** as being a useful tool compound to further probe the biology of ES cells in the context of relocalizing FKBP-EWS/FLI1. For example, future studies could investigate how **EB-TCIP** impacts transcriptional condensate formation, which is an important mechanism for gene activation by EWS/FLI1 and other TFs^38,39^. Potentially, **EB-TCIP** may be forming new transcriptional condensates that contain both EWS/FLI1 and BCL6. Although the utility of **EB-TCIP** may be limited for phenotypic measurements, as we observed off-mechanism cytotoxicity, we have also gained insights that may help inform the next generation of EWS/FLI1 TCIPs, such as their proteasome dependent activity.

Our study is important because it demonstrates that TCIP molecules can bring together two DNA binding proteins. An unforeseen activity of **EB-TCIP** was its ability to induce the degradation of BCL6. We show that proteasome inhibition negates the degradation of BCL6 and impairs the activity of **EB-TCIP**. With global protein degradation inhibited, BCL6 and FKBP-E/F levels increase, which we hypothesized would increase EB-TCIP induced ternary complex and BCL6 target expression. However, we observed the opposite and since proteasome inhibition should not limit transcription, these data suggest that BCL6 degradation is important for maximal activity of **EB-TCIP**. This is also corroborated by our ChIP-seq data, as we generally observed decreases in BCL6 binding at sites where FKBP-E/F binding is gained and chromatin is opened. However, BCL6 degradation alone with **BI3802** treatment is not enough to activate transcription to the same level as **EB-TCIP**. The degradation of BCL6 induced by **EB-TCIP** was competed away with excess **OAP** and not observed when cells were treated with **NEG-2**, which retains the ability to bind BCL6, suggesting that FKBP-E/F or a protein associated with FKBP-E/F, is inducing BCL6 degradation. Therefore, future EWS/FLI1 relocalizing TCIP molecules may also induce degradation of the targeted repressor.

Although our study focused on hijacking EWS/FLI1 TF activity, another interesting avenue of investigation would be recruiting genetic repressors to EWS/FLI1 to decrease oncogenic gene expression. Our chromatin data shows BCL6 moves to FKBP-E/F loci in a **EB-TCIP** dependent manner. Even though BCL6 is redirected to FKBP-E/F sites, some sites show increases in gene expression rather than a decrease. The inability of BCL6 to consistently repress EWS/FLI1 activated genes could be for several reasons. First, FKBP-E/F binding does not change at these sites and the transcriptional activation activity of FKBP-E/F may out compete the repressor activity of BCL6. Second, the magnitude of BCL6 gained at FKBP-E/F sites is lower (>10 fold) than the BCL6 present at *SOCS2* or *CISH*. Therefore, there may not be enough BCL6 gained at these sites to repress gene expression. Finally, **EB-TCIP** uses an inhibitor of BCL6, and the BCL6 that is recruited to EWS/FLI1 may not be functional (e.g., the co-repressor complex is disrupted). TCIPs containing repressor ligands that do not inhibit their function may result in bivalent molecules that can repress expression of EWS/FLI1 activated genes. Alternatively, repressors with stronger repressive function may need to be recruited.

Next generation EWS/FLI1 TCIP molecules will need to address the limitations of **EB-TCIP**. First, endogenous EWS/FLI1 will need to be recruited. Although there is a lack of ligands for EWS/FLI1, future TCIPs could incorporate **MS0621**^40^, a recently described molecule that is reported to interact with EWSR1, EWS/FLI1, and SWI/SNF complex members. Although other molecules, such as **YK-4-279** and its clinical derivative **TK-216**^41^, are reported EWS/FLI1 inhibitors, these molecules may not be ideal ligands for TCIP development as they are also reported to destabilize microtubules at therapeutically relevant concentrations^42^. In light of these shortcomings, our study should encourage EWS/FLI1 ligand discovery as even functionally agnostic compounds could be used to relocalize EWS/FLI1.

Future TCIPs will need to direct EWS/FLI1 to repressors that are ES dependencies. TCIPs targeting BCL6 are antiproliferative in DLBCL because B cell lymphoma cells depend on BCL6 to evade apoptosis. BCL11B, another C2H2 zinc finger repressor, would be a good candidate for next generation EWS/FLI1 TCIPs since it is a known ES dependency^43^. Although there is no reported BCL11B ligand, there is precedence for targeting these proteins with regulatory domain inhibitors (i.e., **BI3812** for BCL6) or iMIDs that bind the C2H2 zinc finger in a cereblon-dependent manner^44^. An interesting repressor candidate in which chemical tools may already exist is ZEB2. Like BCL11B, ZEB2 is a known ES dependency^45^. Recently, it was shown that ZEB2 forms a complex with the lysine demethylase KDM1A in T-ALL cells^46^, which are also dependent on ZEB2. Presumably, KDM1A interacts with ZEB2 at genomic loci were ZEB2 acts as a repressor. There are several classes of known KDM1A inhibitors^47,48^, some with reported anti-proliferative activity in ES^49^, that could be used to recruit EWS/FLI1 to these genomic loci to enhance expression of ZEB2 repressed genes that may be more relevant to ES tumor survival.

The TCIP platform is a promising therapeutic modality for ES and other fusion TF driven cancers. Solid and hematological pediatric malignancies are driven by fusion TFs, such as PAX3/FOXO1^50^ in rhabdomyosarcoma and CBFA2T3/GLIS2^51^ in an aggressive subtype of acute myeloid leukemia (AML). FKBP tagged fusion TF systems could help determine if TCIPs can be used to relocalize fusion TFs beyond EWS/FLI1. ES, rhabdomyosarcoma, and CBFA2T2/GLIS2 AML express fusions that are unique to the tumor cell and are not expressed in healthy cells. Therefore, TCIPs hijacking the fusion TF may have an improved therapeutic window compared to standard chemotherapies/targeted therapies as TCIPs may exhibit reduced toxicity in non-cancerous cells. Building on our proof-of-concept study, future EWS/FLI1 relocalizing TCIPs could serve as novel targeted ES therapies with improved efficacy and safety profiles for patients.

## Acknowledgements

We would like to thank the Molecular Biology Core Facilities Genomics team at DFCI for sequencing our ChIP and ATAC-seq experiments. This work utilized an Illumina NovaSeq X Plus that was purchased with funding from a National Institutes of Health SIG grant 1S10OD036228. NMR spectra were acquired on NMR instruments purchased with funding from the National Insitutes of Health High-End Instrumentation Grant S10OD028697-01 (Stanford ChEM-H). Additionally, this work was funded by support from National Cancer Institute R35 CA283977 and R01 CA283395 (K.S.), Cancer Moonshot U54 CA231637 (K.S. and N.S.G.), National Cancer Institute F32 CA284750 (M.J.B), and National Institute of General Medical Sciences of the National Institutes of Health T32GM136631 (R.P.G). Biorender was used to make images presented in Figure 2A, SI-2B, and SI-3D.

## Materials and Methods

### Data Avaliability

All genome-scale dependency and expression data are available at the DepMap portal website: https://depmap.org. Graph Pad Prism 10 was used to calculate differences between *BCL6* expression in DLBCL vs ES (unpaired T-test with Welch’s correction) and differences in *BCL6* dependency between all other cancers vs DLBCL and all other cancers vs ES (one-way ANOVA). All functional transcriptomics and genomics have been made publically available at the Gene Expression Omnibus (GEO) repository (https://www.ncbi.nlm.nih.gov/geo/) as GSE290895 (RNA-seq), GSE290894 (ChIP-seq), and GSE290893 (ATAC-seq).

### Cell Lines and Reagents

All cell lines used were subject to short tandem repeat (STR) analysis for genotyping and tested for *Mycoplasma* using the MycoAlert® test kit (Lonza, LT07-318). HEK293TF cells used to generate lentivirus were purchased from Thermo Fisher Scientific (#R70007) and grown in Dulbecco’s modified Eagle’s medium (DMEM; Thermo Fisher Scientific, MT10013CM) supplemented with 10% fetal bovine serum (FBS; Sigma Aldrich, F2442) and 1% penicillin-streptomycin (Life Technologies, 15140163). The EWS502 cell line (originally derived in Dr. J. Fletcher’s Lab at Harvard University) was generously provided by Dr. Stephen L. Lessnick of Nationwide Children’s Hospital and all EWS502 lines were grown in Roswell Park Memorial Institute (RPMI)-1640 medium (Life Technologies, 11875119) supplemented with 15% FBS and 1% penicillin-streptomycin. The TC32 cell line (originally derived by Dr. T. Triche at UCLA School of Medicine) was generously provided by Dr. Todd Golub of the Broad Institute and Dana-Farber Cancer Institute (DFCI), and all TC32 lines were grown in RPMI-1640 medium supplemented with 10% FBS and 1% penicillin-streptomycin. To passage cells for maintenance and experiments, cells were washed with sterile phosphate buffered saline (PBS; Life Technologies, 10010023) and detached with 0.05% trypsin-EDTA (Life Technologies, 25300062). Puromycin (Life Technologies, A1113803) and Blasticidin S HCl (Life Technologies, A1113903) were used to select cells as indicated below. Compounds used in this work were acquired from the following sources: **BI3812** (S8735) and **MLN4924** (S7109) were purchased from Selleck Chem. **BI3802** (HY-108705) and **ortho-AP1867** (HY-114434) were purchased from Med Chem Express. **MG132** (474790) and **Actinomycin D** (A4262) were purchased from Sigma-Aldrich.

### Lentiviral CRISPR/Cas9 plasmid construction

Parent plasmids used for guide cloning include lentiCRISPR v2-Blast (Addgene #83480) and lentiCRISPR v2-Puro (Addgene #98290). FastDigest Esp3I (BsmbI; Thermo Scientific, FD0454) was used to digest each backbone, which was then purified by gel extraction (Qiagen, 28704). Synthetic oligonucleotides encoding gene-targeting single guide RNA (sgRNA) sequences (provided below) were purchased from Integrated DNA Technologies (IDT). sgRNAs were annealed and end-phosphorylated using T4 polynucleotide kinase (New England Biolabs, M0201S) in T4 DNA Ligase Reaction Buffer containing 10 mM ATP (New England Biolabs, B0202S). Ligated vectors were transformed into One Shot Stbl3 *Escherichia coli* (Life Technologies, C737303), shaken at 37 °C for 1 h, spread onto 100 µg/mL ampicillin Luria broth (LB) plates (Teknova, L1004) with a L-shaped cell spreader (Fisher Scientific, 14665230) and then grown overnight at 37 °C. Selected colonies were grown overnight in 5 mL of LB (Invitrogen, 12795-027) supplemented with 100 µg/mL ampicillin (Sigma-Aldrich, A9393). Plasmids were DNA-extracted (Qiagen, 27104) and submitted for Sanger sequencing validation at Genewiz (Azenta Life Sciences). Validated clones were cultured overnight in 250 mL volumes, and plasmids were extracted (Zymogen, D4203).

All guides used in this work were from the Broad Institute’s Avana CRISPR-Cas9 library (https://depmap.org). The following guides sequences were used: sgFLI-2 (5’-GATCGTTTGTGCCCCTCCAA-3’), sgBCL6-1 (5’-AGATCCTGAGATCAACCCTG-3’), and sgBCL6-2 (5’-GATCCTGAGATCAACCCTGA-3’). As previously described^52^, sgChr2.2 (5′-GGTGTGCGTATGAAGCAGTG-3′) served as a cutting control and targets a gene desert on chromosome 2. sgLacZ (5′-AACGGCGGATTGACCGTAAT-3′) served as a non-targeting, transduction control and targets a non-human gene. For ligation into the lentiCRISPRv2 (either Blast or Puro) plasmid, the additional bases 5′-CACCG-3′ were added to the 5′ end of the forward sequence. 5′-AAAC-3′ and 5′-C-3′ were added at the 5′ and 3′ ends of the reverse sequence, respectively. sgFLI-2 was cloned into the lentiCRISPR v2-Blast plasmid. sgLacZ, sgChr2.2, sgBCL6-1, and sgBCL6-2 were cloned into the lentiCRISPR v2-Puro vector.

### Generation of polyclonal FKBP-EWS/FLI1 and FKBP-GFP expressing cells

EWS502 cells expressing FKBP-EWS/FLI1 concurrent with knock out of endogenous EWS/FLI1 (sgFLI-Ex9: 5’-GCCTCACGGCGTGCAGGAAG-3′) as well as EWS502 cells expressing FKBP-GFP were generated as described previously^28^. TC32 cells expressing FKBP-EWS/FLI1 were generated in a similar manner, except cells were co-transduced with viral supernatants containing pLEX_305-dTAG-EWS/FLI and lentiCRISPR v2-Blast-sgFLI-2. Cells were then selected and maintained in 1 µg/mL puromycin and 10 µg/mL blasticidin. EWS502-FKBP-EWS/FLI1 cells were also maintained in 1 µg/mL puromycin and 10 µg/mL blasticidin. EWS502-FKBP-GFP cells were maintained in 1 µg/mL puromycin.

### Lentivirus Production and polyclonal CRISPR Cas9 KO of BCL6

CRISPR-Cas9 constructs were packed into lentiviral particles via transduction of HEK293TF cells in Falcon 6 well tissue culture treated plates (Corning, 353046). HEK293TF cells were seeded at a density of 400,000 cells/mL per well. The next day each well was co-transfected with 1250 ng of lentiCRISPR v2-Puro-sgRNA or FgH1tUTG-sgRNA construct plasmid, 250 ng of pVSVG plasmid (Addgene #8454), and 1250 ng of pPAX2 plasmid (Addgene #19319) using Lipofectamine 2000 (Life Technologies, 11668027) according to the manufacturer’s recommended protocol. Plasmids and Lipofectamine 2000 were diluted and mixed in Opti-MEM (Life Technologies, 1058021).

Mixtures of DNA and lipofectamine were added dropwise to each well followed by incubation for 8-16 h at 37 °C, after which media was aspirated and replaced with 3 mL of fresh DMEM. Forty-eight hours after the media change, virus-containing media was collected in 10 mL Luer-Lok syringes (BD, 302995) and sterile-filtered through 0.45 µm syringe filters (Corning, 431225). All infections were performed with freshly produced virus.

For *BCL6* KO experiments, 2 x 10^6^ EWS502 or TC32 cells were seeded into 6-well plates in a volume of 1 mL of RPMI media supplemented with 8 or 4 µg/mL of polybrene (Santa Cruz Biotechnology, SC-134220), respectively. One mL of virus containing media was then added dropwise (final polybrene concentration of 4 or 2 µg/mL) and cells were spin-infected at 30 °C at 2000 rpm for 2 h in a Sorvall Legend XTR centrifuge (Thermo Fisher Scientific). Cells were then incubated at 37 °C overnight. The next day, cells were lifted with trypsin from the 6-well plate and selected with 1 µg/mL of puromycin in a T75 flask (Thermo Fischer Scientific, 156753) for 72 h. Separate samples of non-infected cells subject to the same conditions were treated with puromycin to confirm cell death.

### RNA sequencing (RNA-seq)

For all *BCL6* KO experiments three separate wells of cells were transduced and selected as described above. Approximately two million cells transduced with control or *BCL6* sgRNAs from each well were washed with PBS and then detached from the plate using trypsin. Half of the cells were aliquoted into a 1.5 mL Eppendorf tube and were set aside for protein purification to confirm KO. The other half of cells were pelleted by centrifugation at 2500xg for 3 min in a tabletop centrifuge (Eppendorf, 5425). Media was aspirated and total RNA was extracted using the RNAeasy Plus kit (Qiagen, 74134). Preparation of RNA-seq libraries from total RNA and sequencing was performed by Novogene (https://en.novogene.com). Sequencing was done at ∼20 million reads per sample.

Per Novogene correspondence, RNA integrity was assessed using a Bioanalyzer 2100 System (Agilent Technologies). Libraries were then prepared by purifying messenger RNA (mRNA) from total RNA samples using poly-T oligo-attached magnetic beads. Purified mRNA was fragmented and library prep completed using Fast RNA seq Lib Prep Kit V2 (AbClonal Technology, RK20306). Library quality and concentration were assessed using real-time PCR and Qubit fluorometric quantitation (Thermo Fisher Scientific). Libraries were then pooled based on concentration and sequenced in 150-bp paired-end fashion on a Novaseq6000 instrument (Illumina).

For RNA sequencing experiments of EB-TCIP treated cells, 800,000 EWS502 FKBP-E/F cells were seeded into a 6 well plate. The next day, cells were treated in sextuplicate with DMSO, 2.5 µM **BI3812**, or 2.5 µM **EB-TCIP**, or 2.5 µM **NEG-1**. One set of triplicates was collected as described above 8 h post treatment and the second set of triplicates was collected as described above 24 h post treatment. At both time points cells collected for RNA harvesting were frozen in 350 µL of RLT plus buffer (Qiagen) at -80 °C. All samples were thawed at the same time and total RNA purified using the RNAeasy Plus kit. Total RNA was then subjected to library prep and RNA-seq by Novogene as described above.

### Quantitative-Real Time PCR (qPCR)

Total RNA from 400,000 to 800,000 cells was extracted using the RNAeasy Plus kit. If cells were split for protein and RNA isolation, trypsin was used to detach cells from the plate as described earlier. For experiments where only RNA was harvested, cells were lysed in RLT plus buffer on the plate. Between 1 and 1.5 µg of total RNA was reverse transcribed into cDNA using the High-Capacity cDNA Reverse Transcription Kit (Applied Biosystems, 4368814) and then diluted 1:5 with UltraPure DNase/RNase-Free Distilled Water (Invitrogen, 10977015). All qPCR reactions were performed using the TaqMan system (Thermo Fisher Scientific) with technical triplicate or quadruplicate. Probes used in this study include: *SOCS2*: Hs00919620_m1, *CISH*: Hs00367082_G1, *BCL6*: Hs00153368_m1, and *GAPDH*: Hs02786624_G1 (60x primer limiting). In each qPCR reaction, the gene of interest was measured using FAM dye while the *GAPDH* control was measured using VIC dye. Samples were analyzed in 384-well plate format using 5 µl of either TaqMan Universal Master Mix (Thermo Fisher Scientific, 4304437) or Fast Advanced Master Mix for qPCR (Thermo Fisher Scientific, 4444557), 0.5 µl of FAM-emitting probe, 0.17 µl of VIC-emitting GAPDH probe (60x stock), 2 µl of diluted cDNA and 2.33 µL of UltraPure water for a total of 10 µl per reaction. From the 10 µL reaction volume, 8 µL were pipetted into a MicroAmp Optical 384-well plate (Thermo Fisher Scientific, 4309849) using a 0.5-12.5 µL E1-ClipTip electronic pipet (Thermo Scientific). The plate was spun briefly and then sealed with an optical adhesive cover (Thermo Fisher Scientific, 4360954). The QuantStudio 6 Flex Real-Time PCR machine and the accompanying QuantStudio Real-Time PCR software v.1.7 (Thermo Fisher Scientific) was used to produce and analyze data. The delta-threshold cycle number (ΔCt) was calculated as the difference in threshold cycle number (Ct) between the gene of interest and *GAPDH*. The ΔΔCt was calculated as the difference between the ΔCt of a particular sample and the average ΔCt of the DMSO-treated or sgLacZ control samples. The fold change in gene expression (after *BCL6* KO or compound treatment) was calculated as the ratio of 2^−ΔΔCt^ in sgLacZ cells vs other guides or DMSO treated cells vs cells treated with other compounds. Microsoft Excel was used to calculate ΔCt, ΔΔCt, and fold change in gene expression.

One-way ANOVAs were used to compare changes in gene expression between control conditions (sgLacZ or DMSO). For time course experiments, one-way ANOVAs were used to compare the mean of **EB-TCIP** to all other conditions. For experiments comparing parental, FKBP-GFP, and FKBP-E/F cells, the ratio of **BI3812** induced expression compared to DMSO vs **EB-TCIP** induced expression compared to DMSO was calculated in Microsoft Excel. One-way ANOVA statistics were also used to compare differences between treatments for each cell type. All ANOVA statistics were calculated with Graph Pad Prism 10 using technical replicates.

### Generation of EWS502 FKBP-EWS/FLI1 BCL6 GFP reporter and flow cytometry

The BCL6 GFP reporter plasmid used in previous TCIP publications^8,9^ was graciously provided by the lab of Dr. Jerry Crabtree of Stanford University. Lentiviral particles containing the construct were produced as described above. EWS502-FKBP-EWS/FLI1 cells were infected with the lentiviral particles. Cells were selected for 72 h with 1 and 10 µg/mL of puromycin and blasticidin, respectively. After selection, cells were sorted on a BD Symphony S6 UV Cell Sorter at the DFCI Flow Cytometry Core, which yielded a polyclonal cell population with uniform GFP signal. From this population, single clones were selected by plating 0.5 cells/well into 96 well plates. After four weeks, single colonies were harvested and expanded. The clone that displayed the brightest GFP fluorescence by flow cytometry after 1 µM **EB-TCIP** treatment for 24 h was selected for further experiments.

Fifty-thousand reporter cells were plated per well in a Falcon 24 well-plate (Corning, 353047). The next day cells were treated with a dose response of **EB-TCIP**, **NEG-1**, **NEG-2**, or DMSO. Twenty-four hours later, cells were collected, filtered through Falcon Round-Bottom Polystyrene Test Tubes with Cell Strainer Snap Cap (Fisher Scientific, 0877123) and GFP intensity was measured by flow cytometry at 10,000 cells per sample on a BD FACSCelesta instrument. Live cells were gated using FSC-A and SSC-A. Data was analyzed using FlowJo v.10.4 software. Ratios of the number of cells with GFP intensity >10^3^ in bivalent compound treated cells vs DMSO treated cells were calculated in Microsoft Excel and are reported.

### Time Resolved Fluorescence Energy Transfer (TR-FRET)

Each reaction contained 25 nM His6-TEV-FLAG-FKBP12-F36V, 200 nM BCL6^BTB^-Avi-Biot, 20 nM Streptavidin-FITC (Thermo #SA1001), and 1:400 anti-6xHis terbium antibody (PerkinElmer #61HI2TLF) in 10 uL of buffer containing 20 mM HEPES, 150 mM NaCl, 0.1% BSA, 0.1% NP40, and 1 mM TCEP in a 384-well plate. Protein was incubated with drug digitally dispensed (Tecan D300e) for 1 h in the dark room at room temperature before excitation at 337 nm and measurement of emission at 520 nm (FITC) and 490 nm (terbium) with a PHERAstar FS plate reader (BMG Labtech). The ratio of signal at 520 nm to 490 nm was calculated in Microsoft Excel and normalized to DMSO-treated conditions and plotted.

### Protein Constructs and Purification for TR-FRET

Biotinylated BCL6^BTB^-AviTag protein used for TR-FRET assays included BCL6 amino acids 5-129 with the following mutations: C8Q, C67R, C84N^53^. These enhance stability but do not affect the affinity for BI3812. Preparation of this protein has been described previously^8^.

The construct used for FKBP^F36V^ was pNSG317 (His6-TEV-FLAG-FKBP12-F36V). Rosetta 2(DE3) (Sigma #71400) E. coli cells were transformed with plasmid and inoculated as a starter culture in 50 mL Luria Broth supplemented with chloramphenicol and carbenicillin overnight. Saturated culture was divided into 2L 2XYT medium supplemented with appropriate antibiotics and grown to OD800 = 0.8 at 37 °C. Protein expression was induced by addition of 400 µM IPTG (final concentration, Sigma #I678) and the temperature was adjusted from 37 °C to 18°C for overnight incubation. After incubation overnight, cells were harvested by centrifugation. Cell pellets were resuspended in ∼2 ml/L D800 buffer (20 mM HEPES, pH 7.5; 800 mM NaCl; 10 mM imidazole, pH 8.0; 10 % glycerol, 2 mM beta mercaptoethanol) supplemented with protease inhibitors (1 mM PMSF, 1 mM benzamidine, ∼20 ug/ml pepstatin, aprotinin, and leupeptin) and frozen at -80°C. Cell pellets were thawed briefly in warm water and lysed by sonication and addition of solid egg white lysozyme (Goldbio, L-040-10) before centrifugation at 16,233xg for 1 h at 12°C. Clarified lysate was mixed with ∼0.5 ml/L of growth cobalt resin (Goldbio) for 1h before centrifugation at low speed to separate the beads, which were then washed by gravity flow with ∼25 column volumes ice cold D800 buffer before a final wash with B50 (D800 with 50 mM NaCl) and elution with C50 (B50 with 400 mM imidazole, pH 8.0). Cobalt eluate was applied to a 5 ml anion exchange column (Q HP, Cytiva) and eluted with an 8-column volume gradient from B50 to D800. After concentration in a 3,000 MWCO Amicon filter (Millipore #UFC9003), the sample was applied to a 24 ml gel filtration column (S200 increase, Cytiva) primed with GF150 buffer (20 mM Tris-HCl, pH 8.5, 150 mM NaCl, 1 mM TCEP). S200 peak fractions were again concentrated by ultrafiltration, supplemented with 5% glycerol (*v:v,* final), and aliquoted and frozen at -80°C. A fresh aliquot was thawed for each assay.

### Lysate Preparation and Immunoblotting

Cells were lysed in Radio Immunoprecipitation Assay (RIPA) lysis and extraction buffer (Thermo Scientific, 89900) supplemented with Halt protease inhibitor (Thermo Scientific, 87786) and Halt phosphatase inhibitor (Thermo Scientific, 78420). For on plate lysis, plates with attached cells were placed on ice for 3 min, media aspirated, and then cells were washed with ice-cold PBS. PBS was aspirated and RIPA buffer was added for 15 min with plates on ice. Cells were scraped into chilled 1.5 mL Eppendorf tubes, vortexed for 20 sec and then placed on ice for 15 min, after which the lysate was vortexed for another 20 sec. Lysates were then clarified at 21,100xg for 20 min at 4 °C in a Sorvall Legend Micro 21R centrifuge (Thermo Scientific). For experiments in which protein and RNA were isolated, cells were harvested as described above. Suspended cells were placed on ice for 3 mins and then pelleted at 2500xg at 4 °C. Media was aspirated and the pellets were washed with 1 mL of ice cold PBS followed by another centrifugation at 2500xg at 4 °C. PBS was aspirated and pelleted cells were then resuspended in RIPA buffer, vortexed for 20 secs every 15 min over a 30 min period, and then clarified as described above.

Lysates were prepared for gel electrophoresis by adding 4X NuPAGE LDS loading buffer (Life Technologies, NP0007) supplemented with 10% β-mercaptoethanol (BME, Sigma-Aldrich, M6250). Before addition of loading buffer, protein was quantified by colorimetric Pierce BCA assay (Thermo Scientific, 23227). One microliter of lysate was mixed with 100 µL of BCA:4% copper(iv) sulfate pentahydrate (50:1) in a Falcon 96 well plate (Corning, 353072). The plate was incubated at 37 °C for 30 min and then absorbance read at 562 nm on a Benchmark Plus microplate spectrophotometer (BioRad). The linear correlation from a standard curve of 0, 1, and 5 ug/µL was used to calculate protein concentrations in Microsoft Excel. Thirty to 45 µg of protein was run on 4-15% 1.5 mm NuPAGE Bis-Tris mini pre-cast gels (Thermo Fisher Scientific, NP0336) using NuPAGE MOPS SDS Running Buffer (Thermo Fishcer Scientific, NP0001). Protein was run at 80-90 V for ∼20 min and then run at 130-145 V for an additional ∼90 min. Once electrophoresis was complete, protein was transferred to a 0.2 µm nitrocellulose membrane (BioRad, A30741963) using the Trans-Blot Turbo System (BioRad) at 1.3 A and 25 V for 10 min. Membranes were then incubated in 1X Tris Buffered Saline (TBST; Boston BioProducts, IBB-181) for 3 min with agitation. Next, membranes were blocked for 15 mins at room temperature in EveryBlot Blocking Buffer (EBB; BioRad, 12010020). Membranes were then cut at 25 kDa and 50 kDa and incubated in primary antibody diluted in EBB supplemented with 0.02% sodium azide (Sigma-Aldrich, S2002) overnight (12-16 h) at 4 °C with agitation. The next morning membranes were washed three times with 5 mL of TBST for 5 mins per wash at room temperature. Membranes were then incubated in anti-rabbit IgG HRP-linked secondary antibody (Cell Signaling Technologies (CST), 7074S) diluted 1:10,000 in TBST for 1 h at room temperature. Next, membranes were washed three times with 5 mL of TBST for 5 mins each at room temperature. Protein signal was then visualized using SuperSignal West Femto Maximum Sensitivity Substrate (Thermo Scientific, 34096). Stable peroxide buffer was mixed 1:1 with the luminol/enhancer for 30 sec after which the blot was incubated in the mixture for 1 min before visualizing on a ChemiDoc MP Imaging System (BioRad, 10000062126) using 2×2 binning with rapid or optimal automated exposure. When probing for EWS/FLI1 after BCL6, blots were stripped using Restore Western Stripping Buffer (Life Technologies, 21059) for 1h at room temperature. Blots were washed three times with TBST for 5 mins each at room temperature and then reblocked for 15 mins with EBB. EWS/FLI1 primary antibody diluted in EBB was then added, incubated overnight at 4 °C and imaged as described above. Image Lab Version 6.1.0 build 7 was used to export image files for figures.

The following primary antibodies were used at the following dilutions: rabbit monoclonal anti-SOCS2 (Abcam, ab109245) at 1:1000, rabbit monoclonal anti-BCL6 (CST, 14895) at 1:1000, rabbit monoclonal anti-FLI1 (Abcam, ab133485) at 1:1000, rabbit monoclonal anti-HA (CST, 3724) at 1:1000, and rabbit monoclonal anti-GAPDH (at 1:2000).

### Ternary Complex Pulldowns

EWS502 FKBP-EWS/FLI1 cells growing on 15 cm^2^ dishes (Thermo Scientific, 150350) were washed with PBS, lifted with trypsin, trypsin neutralized with RPMI media, and then pelleted at 1400 RPM for 3 mins in an Eppendorf 5910 R centrifuge. Trypsin/media was aspirated, and the cells were washed with 5 mL of PBS and then counted using a Countess 3 cell counter (Invitrogen). Cells were then pelleted again at 1400 RPM for 3 mins, PBS was aspirated and the cells placed on ice for 5 mins. Next, cells were resuspended in IP lysis buffer (20 mM Tris pH 7.5 (diluted from 1M Tris pH 8.0, Invitrogen, AM9856), 150 mM NaCl (diluted from 5M, Invitrogen, AM9759), and 1% NP-40 (diluted from 10%, Abcam, ab142227)) supplemented with Halt protease and phosphatase inhibitors at a concentration of 10 x 10^6^ cells per 250 µL of lysis buffer. Lysate was kept on ice and vortexed for 20 sec every 15 mins for 1h. Lysate was transferred to a chilled 1.5 mL Eppendorf tubes and cleared at 21,100xg for 20 mins at 4 °C. Lysate was pooled into one chilled 15 mL Falcon tube and then split into 250 µL aliquots in separate, chilled tubes. Twenty-one microliters of lysate were saved as the input sample and mixed with 7 µL of 4X LDS buffer supplemented with 10% BME. Each tube of lysate was then treated with either 0.25 µL of DMSO or 1000x stock of the indicated compound. Lysate was incubated with compounds for 1 h at 4 °C with agitation. For competition experiments, lysates were pretreated with 1000x stocks of **BI3812** or **OAP** or 0.25 µL DMSO for 1 h before addition of **EB-TCIP.** While lysates incubated with compound, 25 µL of Pierce Anti-HA Magnetic Beads (Thermo Scientific, 88837) per pulldown was aliquoted into a 1.5 mL Eppendorf tube. One milliliter of IP lysis buffer was added and then the tube was placed into a DynaMag-2 magnetic rack (Invitrogen, 12321D) until the solution was clear. Buffer was removed and the beads were washed twice more with 1 mL of IP lysis buffer. After the final wash, the beads were resuspended in 26 µL of IP lysis buffer per pulldown and placed on ice. After the incubation with compounds, 25 µL of washed beads was added to each tube. The beads were incubated with treated lysates overnight (16-24 h) at 4 °C. The next day, samples were quickly spun in a microcentrifuge and then beads separated using the magnetic rack. Beads were washed three times with ice-cold IP wash buffer (20 mM Tris pH 7.5, 150 mM NaCl, 0.01% NP-40), with quick spins in between each wash to remove liquid from the cap of the tube. After the third wash the beads were resuspended in 1.5X LDS Buffer supplemented with 2.5% BME and boiled for 10 mins at 95 °C. Boiled samples were spun at max speed in a tabletop centrifuge for 1 min to collect condensation and then placed on a magnetic rack. Supernatant was loaded into a 4-15% 1.5 mm NuPAGE Bis-Tris mini pre-cast gel and subject to electrophoresis and immunoblotting as described above.

### Time Courses

Seven-hundred thousand EWS502 FKBP-EWS/FLI1 cells were plated into each well of four 6 well tissue culture plates. The next day, wells were treated in sextuplicate with either DMSO, 1 µM **BI3812**, 1 µM **EB-TCIP**, or 1 µM **BI3802**. Cells were then harvested at each time point by trypsinization as described above. At each time point half the cells were collected for RNA extraction and the other half used for protein isolation. RNA samples were frozen at -80 °C in RLT plus buffer while protein samples were frozen at -80 °C in RIPA buffer. All RNA or protein samples were thawed at the same time and processed together in a single batch. Purified RNA was subject to RT-qPCR as described above. Lysates were subject to immunoblotting as described above.

### Competition Assay

One million, two hundred thousand EWS502 FKBP-EWS/FL1 cells were plated into each well of two 6 well tissue culture plates. The next day, cells were treated with either DMSO or 25 µM **OAP** (free acid) for 1h at 37 °C. After this pretreatment, media was aspirated and cells were treated with either DMSO, 25 µM **OAP**, 1 µM **BI3812**, 1 µM **EB-TCIP**, 25 µM **OAP** plus 1 µM **EB-TCIP**, or 1 µM **BI3812** plus 1 µM **OAP** for an additional 4 h at 37 °C. Cells were then harvested by trypsinization as described above. Half the cells were collected for RNA extraction and the other half used for protein isolation. RNA samples were frozen at -80°C in RLT plus buffer while protein samples were frozen at -80 °C in RIPA buffer. All RNA or protein samples were thawed at the same time and processed together. Purified RNA was subject to RT-qPCR as described above. Lysates were subject to immunoblotting as described above.

### Ubiquitin/Proteasome & Transcription Inhibitor Treatment

One million, two hundred thousand EWS502 FKBP-EWS/FL1 cells were plated into each well of three 6 well tissue culture plates. The next day, cells were treated with DMSO, 1 µM **MG132**, 1 µM **MLN4924**, or 1 µM **Actinomycin D** for 1 h at 37 °C. After pre-treatment, media was aspirated and cells were treated with either DMSO, 1 µM **EB-TCIP**, or 1 µM **BI3802** plus and minus each inhibitor for an additional 4 h at 37 °C. Cells were then harvested by trypsinization. Half the cells were collected for RNA extraction and the other half used for protein isolation. RNA samples were frozen at -80 °C in RLT plus buffer while protein samples were frozen at -80 °C in RIPA buffer. All RNA or protein samples were thawed at the same time and processed together. Purified RNA was subject to RT-qPCR as described above. Lysates were subject to immunoblotting as described above.

### Chromatin-Immunoprecipitation sequencing (ChIP-seq)

Eleven million EWS502 FKBP-EWS/FLI1 cells were plated into 15 cm^2^ dishes. The next day cells were treated with DMSO, 1 µM **BI3812**, 1 µM **EB-TCIP**, or 1 µM **BI3802** in quadruplicate. After 24 h, media was aspirated, and cells were harvested by trypsinization as described above. Cells from two 15 cm^2^ plates treated with the same condition were pooled and counted. Forty million EWS502 FKBP-EWS/FLI1 cells per condition (20 million cells per ChIP reaction) were collected in a 50 mL Falcon tube. Cells were pelleted at 300xg for 5 mins and then washed twice in 5 ml PBS. Cells were then crosslinked by resuspension in 10 mL PBS containing 1% methanol-free formaldehyde (Thermo Fisher Scientific, 28906) and rotated slowly by hand for 10 mins at room temperature. The reaction was quenched by addition of 1 mL of 2.5 M glycine (Sigma Aldrich, G7126). Cells were pelleted at 800xg for 5 mins at 4 °C pellets and then washed twice with 10 mL PBS at room temperature supplemented with 1 mM PMSF. After resuspending in the second wash, the cell suspension was split into two chilled 50 mL Falcon tubes (5 mL each). After spinning at 800xg for 5 mins and removing the second PBS wash, cell pellets were flash frozen in liquid nitrogen. When processing samples one set of tubes for all conditions was thawed on ice and a pulldown for either HA or BCL6 was performed as described below.

For each immunoprecipitation (IP), 100 µl of protein A Dynabeads (Thermo Fisher Scientific, 10002D) was washed three times in 1 ml BSA blocking solution (0.5% w/v sterile-filtered BSA in UltraPure H_2_O) and resuspended in 250 µl BAS blocking solution. Beads were pooled and then 10 µg of either anti-HA (Cell Signaling Technologies, 86124SF) or anti-BCL6 antibody (Thermo Fischer Scientific, PA5-27390) per IP was added. Two micrograms of spike-in antibody recognizing a Drosophila-specific histone variant was added (Active Motif, 61686) to normalize samples. The following morning, the antibody-conjugated beads were washed four times in 1 ml BSA blocking solution and then resuspended in 100 µl of the solution per IP and stored at 4 °C.

Frozen, crosslinked cells were thawed briefly on ice and then resuspended in 1 ml of SDS lysis buffer (0.5% SDS, 5 mM EDTA, 50 mM Tris-HCl pH 8.0, 100 mM NaCl, and 0.2% sodium azide) supplemented with Halt protease inhibitor and incubated at room temperature for 2 min with gentle agitation. Lysates were transferred to microcentrifuge tubes and centrifuged at 15,000xg for 10 mins at 4 °C. The nuclear pellet was re-suspended in 950 µl of ChIP IP buffer (2 parts SDS lysis buffer and 1 part Triton dilution buffer, which was composed of 100 mM Tris-HCl pH 8.0, 100 mM NaCl, 5 mM EDTA, 0.2% NaN3 and 5% Triton X-100) supplemented with Halt protease inhibitor. Nine-hundred microliters was then transferred to a milliTUBE (Covaris, 520130). Sonication was performed on an E220 Focus Ultra sonicator (Covaris) at 5% duty cycle, 140 W peak power, 200 cycles per burst, at 4 °C for 25 mins per milliTUBE. Sheared chromatin was transferred to a 1.5 ml tube and centrifuged at 15,000xg for 10 mins at 4 °C. The supernatant of sheared chromatin was transferred to a new reaction tube. To prepare the ChIP DNA input sample, 5 µl of sheared chromatin was transferred to a PCR strip-tube (USA Scientific, 1402-4700) and mixed with 40 µl de-crosslinking buffer (100 mM NaHCO3 and 1% SDS buffer), 1 µl RNAse A (Thermo Fisher Scientific, 12091021) and 1 µl proteinase K (Thermo Fisher Scientific, AM2546). The tube was incubated for 2 h at 65 °C in a ProFlex PCR thermal cycler (Applied Biosystems) to de-crosslink DNA–protein covalent bonds. DNA was isolated using Agencourt AMPure XP bead-based purification at a 1.2 times ratio (Beckman Coulter, A63881). Briefly, beads were mixed with the sample in the PCR tube and incubated for 10 mins at room temperate. Tubes were then placed in a magnetic separation rack (EpiCypher, 10-0008) and washed twice with 500 µL of 80% ethanol. DNA was then eluted in 50 µl Tris-EDTA (SigmaAldrich, 93283) and stored at −20 °C. To the remainder of sheared chromatin was added 100 µL of conjugated bead–antibody solution was. Before addition of the antibody bound beads, 40 ng per reaction of *Drosophila* spike-in chromatin (ActiveMotif, 53083) was added to the pooled antibody bound beads. IP reactions were rotated overnight at 4 °C.

The following day, ChIP reactions were washed twice in 1 ml low-salt buffer (0.1% SDS, 1% Triton X-100, 2 mM EDTA, 20 mM Tris-HCl pH 8.0, and 150 mM NaCl), high-salt buffer (0.1% SDS, 1% Triton X-100, 2 mM EDTA, 20 mM Tris-HCl pH 8.0, and 500 mM NaCl), lithium chloride buffer (0.25 M LiCl, 1% IGEPAL-CH 630, 1% sodium deoxycholate, 10 mM Tris-HCl pH 8.0, and 1 mM EDTA) and then once in 700 µl ice-cold Tris-EDTA buffer (Sigma Aldrich, 93283). Chromatin was eluted using 100 µl fresh ChIP elution buffer (1% SDS and 0.1 M NaHCO3) and rotated at room temperature for 15 mins. Eluate was transferred to PCR tubes and mixed with 8 µl 2.5 M NaCl, 1 µl RNAse A and 1 µl proteinase K. Samples were de-crosslinked for 12–16 h at 65 °C in a thermal cycler. ChIP DNA was extracted from the de-crosslinked samples using AMPure XP beads at a 1.2× ratio as described above and eluted in 20 µl of Tris-EDTA. DNA was quantified using a Qubit dsDNA high sensitivity assay (Q32851). DNA fragment sizes were measured with a Tapestation 2200 instrument (Agilent, ScreenTape, 5067-5584; reagents, 5067-5585).

ChIP-seq libraries were prepared using a NEBNext Ultra II DNA Library Kit for Illumina sequencing (NEB, E7645S) and NEBNext Multiplex Oligos for Illumina sequencing (NEB, E6440S). HA and BCL6 samples were PCR-amplified for 12 cycles. Library pooling and indexing was evaluated with shallow sequencing on an Illumina MiSeq. Subsequently, libraries were sequenced on an Illumina NovaSeq X Plus targeting roughly 40 million, 150bp read pairs per sample by the Molecular Biology Core Facilities at Dana-Farber Cancer Institute.

### Assay for Transposase-accessible chromatin with sequencing (ATAC-seq)

Three-hundred thousand EWS502 FKBP-EWS/FLI1 cells were plated into each well of a 12-well tissue culture plate. The next day cells were treated with DMSO or 1 µM **EB-TCIP** in duplicate. After 24 h, cells were harvested by trypsinization as described above and counted. Next, 100,000 cells from each sample were used to prepare libraries for ATAC-seq using a commercially available kit (ActiveMotif, 53150). The molarity of each library was calculated using a Qubit dsDNA Broad Range Assay kit (Thermo Fisher Scientific, Q32850) and an Agilent TapeStation 2200. Library pooling and indexing was evaluated with shallow sequencing on an Illumina MiSeq. Subsequently, libraries were sequenced on an Illumina NovaSeq X Plus targeting roughly 20 million, 150bp read pairs per sample by the Molecular Biology Core Facilities at Dana-Farber Cancer Institute.

### Cell Viability

Fifty microliters of a 10,000 cell/mL suspension of EWS502 or EWS502 FKBP-EWS/FLI1 were seeded into each well of a white polystyrene 384 well cell culture plates (Corning, 3570). The next day cells were treated with DMSO, **OAP**, **BI3812**, or **EB-TCIP** using a HP D300e Digital Dispenser. Cells were treated with 8-point dose responses starting at 10 µM with 1:2 dilutions. Treated cells were incubated for 72 h at 37 °C, after which 10 µL of Cell-Titer-Glo (Promega, G7573) was added to each well using a 2-125 µL E1-ClipTip electronic pipet (Thermo Scientific). The plate was then incubated at room temperature for 15 mins with 350 rpm rotation in an Eppendorf MixMate. Luminesce was determined using a CLARIOStar Plus plate reader (BMG Labtech). The ratio of between luminescence of compound treated samples to DMSO treated samples was calculated in Microsoft Excel. Dose response curves were then generated by fitting the data to an [inhibitor] vs. dose response non-linear regression using GraphPad Prism 10.

### RNA-seq data analysis

RNA-seq data analysis was performed according to the ENCODE standards (https://www.encodeproject.org/data-standards/rna-seq/long-rnas/). Quality check of unaligned reads was performed using FastQC v.0.11.9 (https://www.bioinformatics.babraham.ac.uk/projects/fastqc/) and MultiQC v.1.14^54^ respectively.

Using STAR v.2.7.11a^55^ the paired end reads were aligned to hg38/gencodev30 with standard parameters –outSAMtype BAM SortedByCoordinate --outSAMunmapped None --outSAMattriubtes NH HI NM MD AS XS --outReadsUnmapped FastX --outSAMstrandField intronMotif --quantMode TranscriptomeSAM GeneCounts --qantTranscriptomeBan IndelSoftclipSingleend --readFilesCommand zcat. Gene level reads were counted and summarized across hg38 exons by using featureCounts v.2.0.3 from the Subread v2.0.0 package (https://subread.sourceforge.net/). Following alignment, quality control checks were performed using SARTools v.1.7.3^56^. DESeq2 v.1.44.0 was used to normalize gene counts and quantify differential expression between experimental and control conditions^57^ using the apeglm v1.26.1^57^ library. Gene level expression was estimated as log_2_(TPM +1) normalized reads. Expressed genes were identified as genes with maximum log_2_(TPM +1) expression ≥ 1 across conditions. Gene differential expression status (decrease, increase or not significant change) was estimated based on shrunken log_2_ fold change scores with the cutoffs |fold change expression| ≥ 1.5 and adjusted P ≤ 0.10. Heatmaps displaying transcriptional changes were created using the Morpheus software platform (https://software.broadinstitute.org/morpheus/) based on log_2_(fold change) expression data.

### Gene set enrichment analysis

GSEA software v.4.2.2^23^ was used to identify signature enrichment of experimental conditions in *BCL6* KO, compound treatment, and corresponding conditions. MSigDB v7.4 collections, a published BCL6 target gene set^24^, and in-house curated gene sets were analyzed for enrichment against the data. For each experimental condition, the expressed genes were ranked based on the expression fold change in sgBCL6 vs sgChr2.2 or compound treated versus DMSO control. Results were visualized with volcano plots with Normalized Enrichment Score (NES) versus -log_10_(P) and GSEA plots. Significance cutoffs for GSEA enrichments: |NES| ≥ 1.3, P ≤ 0.10, FDR ≤ 0.25.

### ChIP-Seq data analysis

The analysis of the spiked-in ChIP-Seq data was performed according to the ENCODE standards (https://www.encodeproject.org/chip-seq/). Quality control was performed on unmapped sequences using FastQC v.0.11.9 (https://www.bioinformatics.babraham.ac.uk/projects/fastqc/) and MultiQC v.1.14^54^. Adapters and low-quality reads were removed using Trimmomatic v0.39^58^. Reads were mapped to the human genome (GRCh38/hg38) and to the spike-in *Drosophila melanogaster* (dm6) using bowtie2 v.2.5.1^59^ with the “local very_sensitive” parameters. Mapped reads were processed with SAMtools v0.1.19^60^ and reads with low mapping quality (MAPQ < 5) were disregarded. Duplicate reads were removed using the Picard Mark Duplicates method implemented in the sambamba 0.7.1 tool^61^. Fragment size distributions were computed using the PEFragmentSize tool available in the deepTools v.3.5.1 package^62^.

Human and *Drosophila* genome-wide counts across 2000 bp bins were computed with the bamSummary tool available in the deepTools package v.3.5.0^62^. Bins with at least 10 reads in less than 3 samples and bins overlapping ENCODE blacklisted regions were excluded. The Active Motif Spike-in Normalization protocol was then applied to compute the scaling factors per antibody samples as ratios between the average dm6 counts across antibody samples vs. the dm6 counts for that sample. The normalization factor was set to 1 if the percentage of *Drosophila* reads was less than the 1% minimum cutoff.

The bamCoverage tool from the deepTools package v.3.5.0^62^ was used to generate normalized reads per kilobase per million (RPKM) genome-wide coverage bigwig files with specified bin sizes of 20 bp and scaled with the pre-computed spike-in scale factors.

Peak calling was performed using the model-based MACS2 v.2.1.1.20160309^63^ software against experimental inputs with a significance cutoff FDR ≤ 0.01. Bwtool software^64^ was used to compute the area under the curve (AUC) for the RPKM normalized signal across genomic regions. MACS2 peaks were filtered by removing binding regions with low AUC coverage of [log_2_(AUC+1)<14] and ENCODE hg38 black-list regions (https://www.encodeproject.org/annotations/ENCSR636HFF/).

Various mapping and genomic analyses including indexing, sorting, intersection, and merging were executed using SAMtools v.1.9 and Bedtools v.2.29^65,66^. Next, quality control for peaks called was performed using ChIPQC^67^ under the Bioconductor package v.3.9. Homer v4.11^68^ platform was used to annotate peaks called with the closest hg38 genes using the annotatePeaks function. Peak binding signal were visualized using the Integrative Genomic Viewer (IGV) v.2.12.3^69^. Promoter regions were defined as the area of the genome ±3.0kb from gene transcription start sites (TSS).

Antibody binding sites identified by MACS2 were merged into a set of aggregated peaks for control and treatments across conditions. Utilizing deepTools multiBamSummary tool, peak by sample counts was generated. Counts were used to perform differential peak analysis. Changes between two conditions binding signal were identified as increase, decrease or not significant based on absolute cutoff of 1.5 for delta area under curve. Significance of changes for binding reads was calculated by using DESeq2 v.1.44.0 with a cutoff of P ≤ 0.10.

Heatmaps of normalized AUC signal were created using deepTools v.3.5.1 computeMatrix and plotHeatmap functions. Metaplots displaying average normalized scores across genomic regions were created using deepTools v.3.5.1 plotProfile function. Motif enrichment analysis was performed using Homer v.4.11^68^.

Box plots of Supplemental Figure SI-6 D-G used Homer annotatePeak genes to map peaks in increasing, decreasing, and unchanged groups to genes from DESeq2 analysis shrunken log2 fold change values. HA peak groups in Supplemental Figure SI-6 D-E were subdivided using bedtools by whether they overlapped with the merged AUC filtered BCL6 peaks. BCL6 peak groups in Supplemental Figure SI-6 F-G were subdivided using bedtools by whether they overlapped with the merged AUC filtered HA peaks. Genes with peaks from the increase or decrease groups were excluded from the no change group. Using the R pairwise.t.test function, a paired t-test with a Benjamini-Hochberg correction was used to evaluate the significance for the differences of the RNA-seq LFC distributions for the various peak categories.

### ATAC-Seq data analysis

ATAC-Seq analysis was performed according to the ENCODE standards (https://www.encodeproject.org/atac-seq/). Quality control was performed on the unmapped paired end reads with FastQC v.0.11.9 (https://www.bioinformatics.babraham.ac.uk/projects/fastqc/) and MultiQC v.1.14^54^. Adapters were then trimmed and filtered using Trimmomatic v.0.36^58^. Using bowtie2 v.2.5.1, the trimmed paired end reads were aligned to the hg38 genome with the -local - very_sensitive -X 2000 parameters. Reads mapped to the hg38 genome to chromosomes 1 to 22 with a MAPQ > 5 were kept. Duplicates were removed using Picard Mark Duplicates method implemented in the sambamba 0.7.1 tool^61^. deepTools v.3.5.1^62^ AlignemntSieve tool was used to shift reads 4bp on the positive strand and -5bp on the negative strand. Replicate correlations were calculated and visualized using multiBamsummary and bamCorrelate, along with fragment size distributions using PEFragmentSize within the deepTools v.3.5.1 package^62^. Replicates were merged and then peak calling was performed with MACS2 v.2.1.1.20160309^63^. Next, AUC binding signal was computed with the bwtool program^64^. The Homer v.4.11^68^ program was employed to annotate called peaks to the closest hg38 genes using the annotatePeaks function. Promoter regions were defined as the area of the genome ±3.0kb from gene transcription start sites (TSS). Peak binding signal was visualized using the Integrative Genomic Viewer (IGV) v.2.12.3^69^.

