## Supporting Information for "Rewiring the fusion oncoprotein EWS/FLI1 in Ewing sarcoma with bivalent small molecules"

23 Pages

Supporting Information Figures 1-8

Chemical Synthesis

Supporting Information Tables 1-3

Supporting Information References

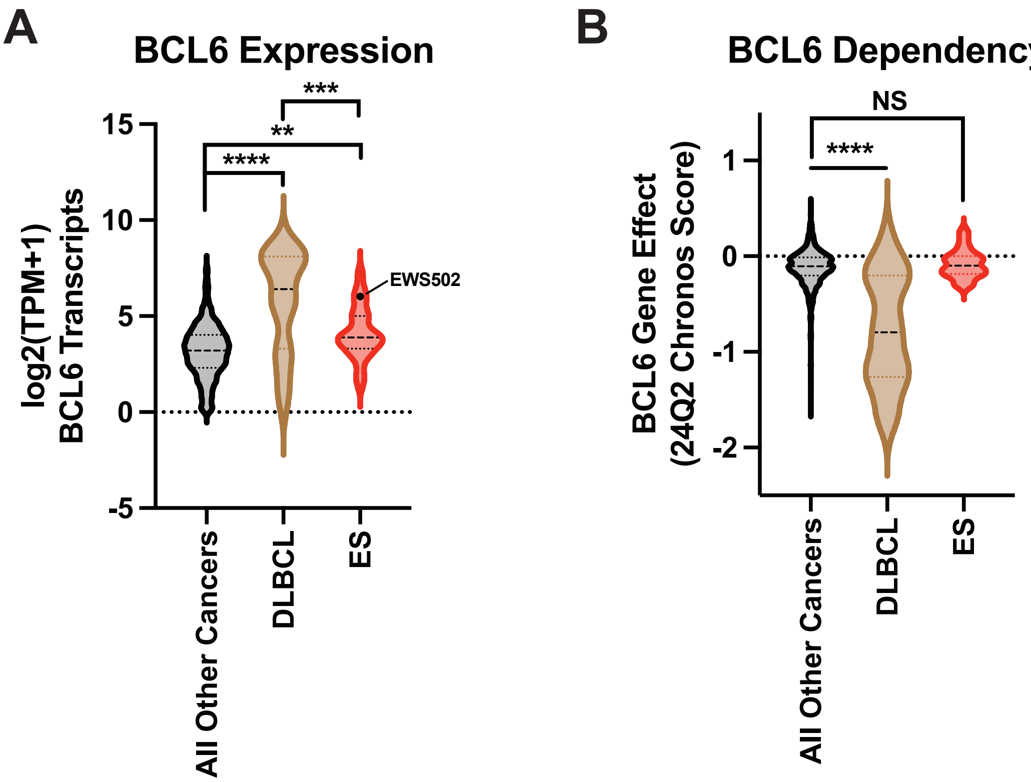


**Figure SI-1: *BCL6* is highly expressed in ES cells, but *BCL6* is not a dependency in ES cells.** (A) comparison of expression ($\log_2$ TPM+1) from DepMap portal of *BCL6* in all other cancers (black) vs DLBCL (brown) vs ES (red). The dot in ES corresponds to expression of EWS502 cells used in this work, which have expression near the $50^{th}$ percentile of DLBCL cells. TC32 *BCL6* expression is not available in DepMap. Means of each group compared with one-way ANOVA; ** $p < 0.01$, *** $p < 0.005$, **** $p < 0.001$. (B) Gene effect (Chronos score) from DepMap portal for *BCL6* in all other cancers vs DLBCL vs ES. The more negative the mean the more dependent the cell type is on *BCL6*. DLBCL is dependent compared to all cancers while ES is not. Means of each group compared with one-way ANOVA; NS = not significant, **** $p < 0.001$.

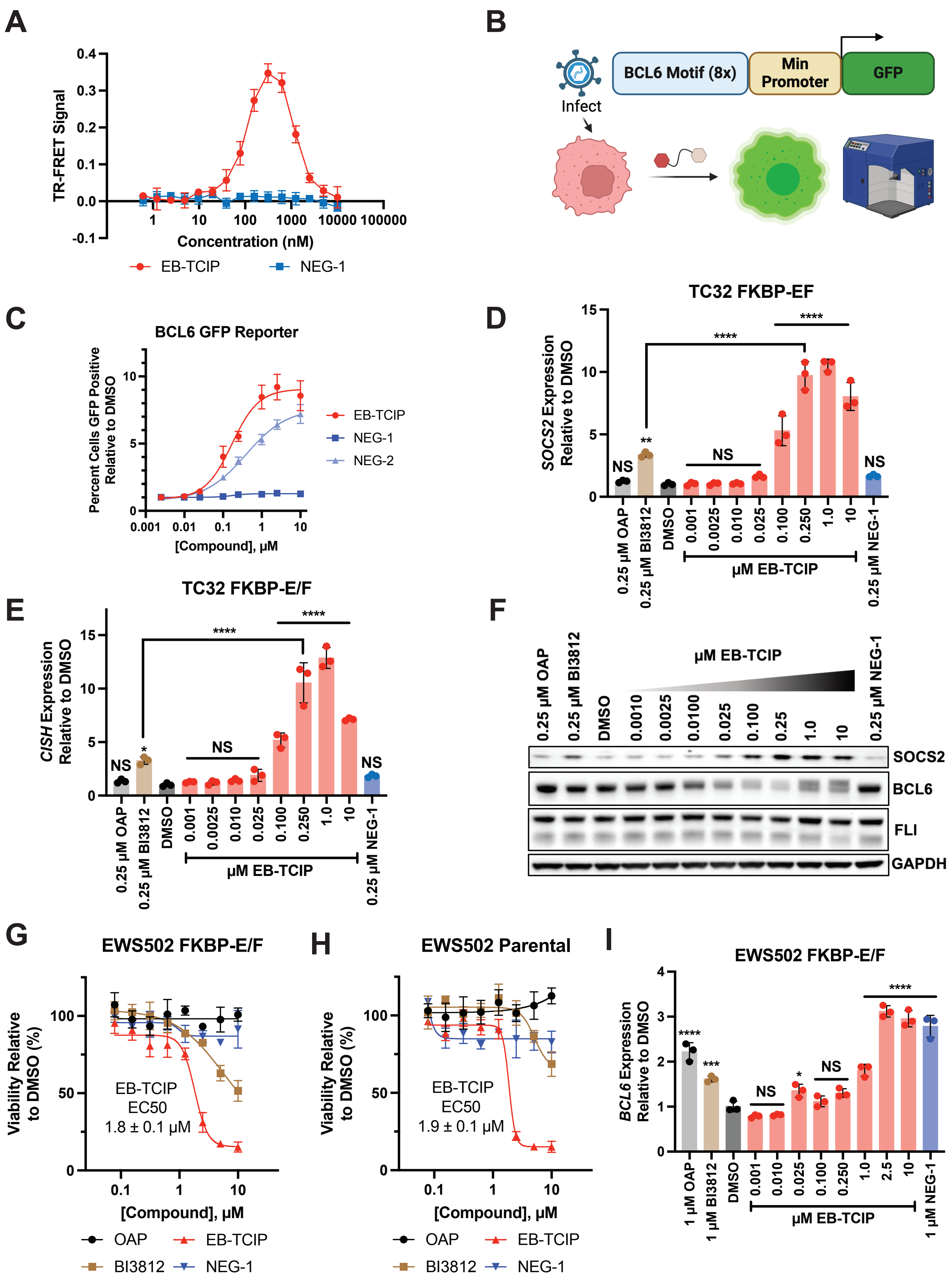


**Figure SI-2: *In vitro* and *in cellulo* characterization of EB-TCIP.** (A) TR-FRET experiment showing **EB-TCIP** (red), but not **NEG-1** (blue), can form a ternary complex between purified FKBP$^{F36V}$ and

BCL6$^{BTB}$. Data shown is one biological replicate, in technical triplicate, that is representative of three biological replicates. Error bars indicate mean ± SD. (B) Schematic of BCL6 GFP reporter assay. Image made in Biorender. (C) Dose response of **EB-TCIP** (red), **NEG-1** (dark blue), and **NEG-2** (light blue) in the FKBP-E/F EWS502 cells expressing the BCL6 GFP Reporter assay. Data shown is the average of three biological replicates. Error bars indicate mean ± SD. **EB-TCIP** dose dependently increases *SOCS2* (D) and *CISH* (E) in by RT-qPCR in TC32 cells expressing FKBP-E/F. (F) SOCS2 protein levels dose dependently increase, while BCL6 protein levels dose dependently decrease in FKBP-E/F TC32 cells. GAPDH was used as a loading control. Blot is representative of three biological replicates. **EB-TCIP** is equally antiproliferative in both EWS502 FKBP-E/F cells (G) and parental EWS502 cells (H). CTG data shown is one experiment performed in technical triplicate that is representative of two biological replicates. Error bars indicate means ± SD. (I) **EB-TCIP** dose dependently increases *BCL6* transcript levels in EWS502 FKBP-E/F cells by RT-qPCR. *BCL6* levels were also elevated by **OAP**, **BI3812**, and **NEG-1**. All RT-qPCR experiments are single experiments, done in technical triplicate, that are representative of two biological replicates. Means were compared by one-way ANOVA; NS = not significant, * $p < 0.05$, ** $p < 0.01$, *** $p < 0.005$, **** $p < 0.001$.

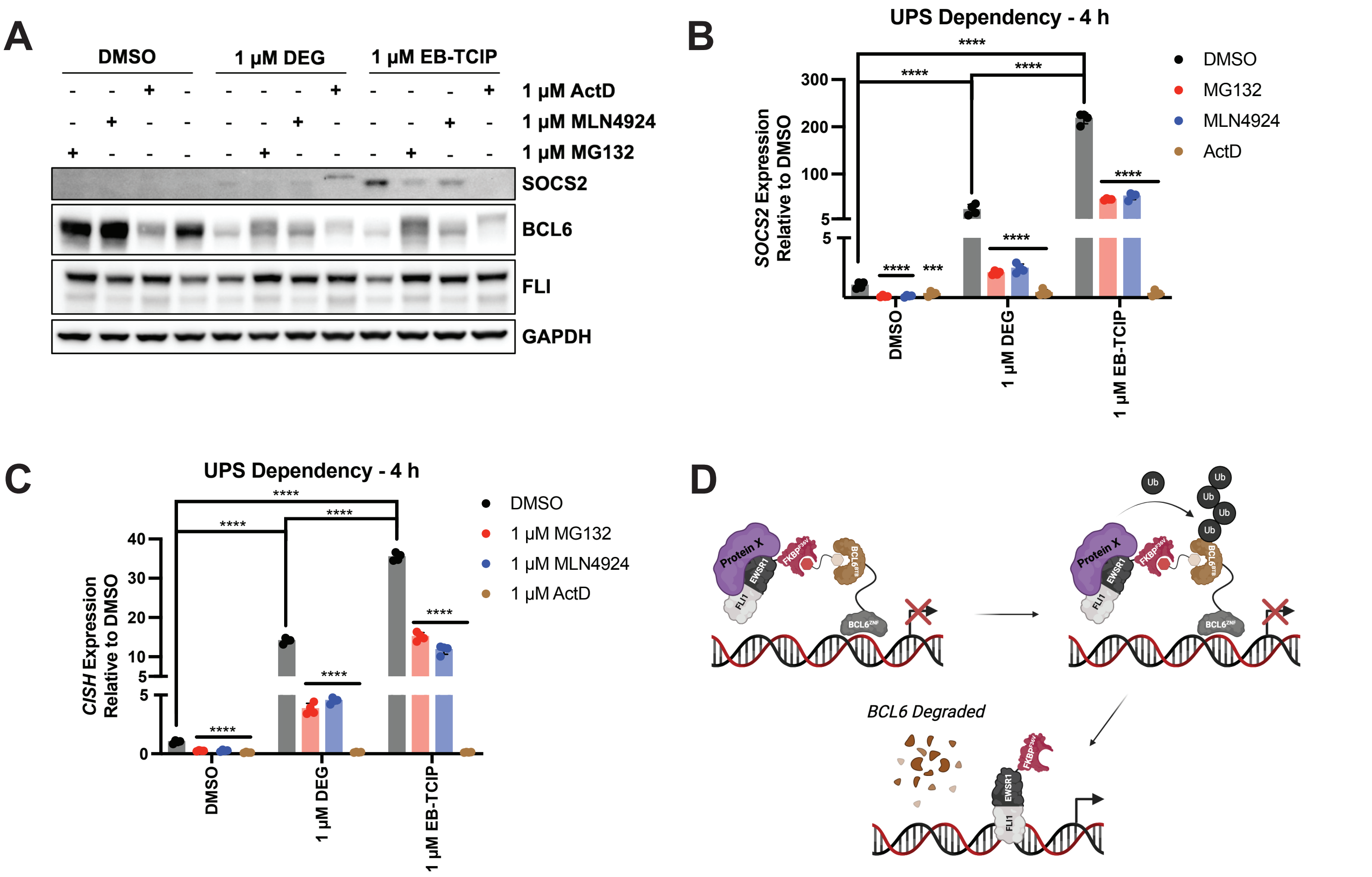


**Figure SI-3: EB-TCIP activity is proteasome dependent**. (A) Pre-treatment of EWS502 FKBP-E/F cells with **MG132**, **MLN4924**, or **Actinomycin D** (**ActD**) (1 μM) inhibits the ability of **EB-TCIP** to increase SOCS2 protein levels, even though BCL6 and FKBP-E/F levels increase with **MG132** and **MLN4924** treatment. **BI3802** (**DEG**) induced degradation of BCL6 and increases in SOCS2 protein levels are also rescued by **MG132** and, to a lesser extent, **MLN4924**. Blot is representative of three biological replicates. Ubiquitin-proteasome (UPS) inhibitor and transcription inhibitor pretreatment decrease expression of *SOCS2* (B) and *CISH* (C) in EWS502 FKBP-E/F cells treated with DMSO, **EB-TCIP**, or **BI3802** (1 μM). All RT-qPCR experiments are a single experiment, done in technical triplicate, that is representative of two biological replicates. Means were compared by one-way ANOVA; NS = not significant, * $p < 0.05$, ** $p < 0.01$, *** $p < 0.005$, **** $p < 0.001$. Unless indicated with brackets, the means of **MG132**, **MLN4924**, and **ActD** treated cells were compared to DMSO for each treatment. (D) Schematic showing how TCIP activity could be UPS dependent. **EB-TCIP** may recruit a protein that induces the degradation of BCL6, opening up sites on chromatin for FKBP-E/F to bind and drive gene expression. Image made in Biorender.

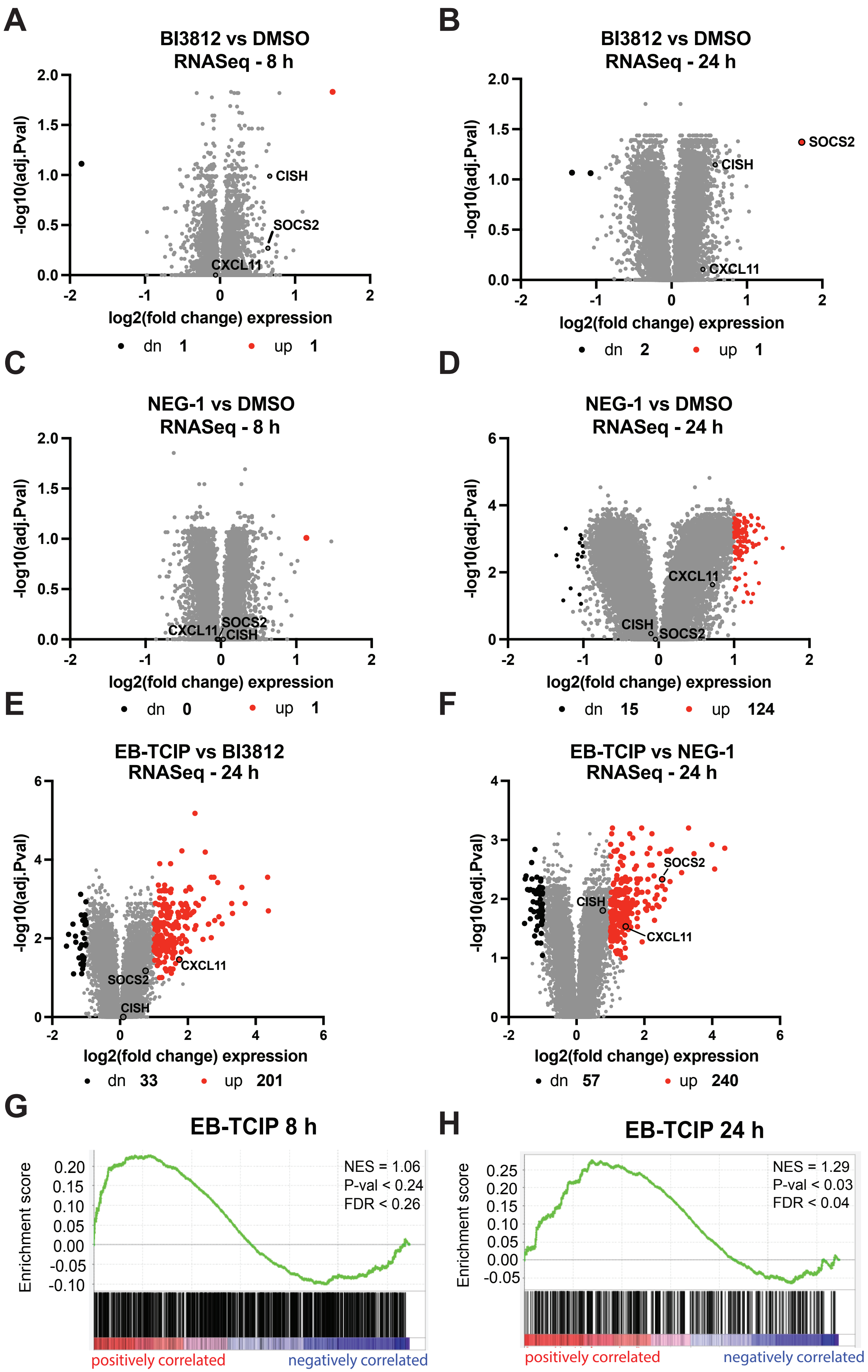
A
BI3812 vs DMSO
RNASeq - 8 h
-log10(adj.Pval)
log2(fold change) expression
CISH
SOCS2
CXCL11
dn 1
up 1
B
BI3812 vs DMSO
RNASeq - 24 h
-log10(adj.Pval)
log2(fold change) expression
SOCS2
CISH
CXCL11
dn 2
up 1
C
NEG-1 vs DMSO
RNASeq - 8 h
-log10(adj.Pval)
log2(fold change) expression
SOCS2
CXCL11
CISH
dn 0
up 1
D
NEG-1 vs DMSO
RNASeq - 24 h
-log10(adj.Pval)
log2(fold change) expression
CXCL11
CISH
SOCS2
dn 15
up 124
E
EB-TCIP vs BI3812
RNASeq - 24 h
-log10(adj.Pval)
log2(fold change) expression
CXCL11
SOCS2
CISH
dn 33
up 201
F
EB-TCIP vs NEG-1
RNASeq - 24 h
-log10(adj.Pval)
log2(fold change) expression
SOCS2
CISH
CXCL11
dn 57
up 240
G
EB-TCIP 8 h
Enrichment score
NES = 1.06
P-val < 0.24
FDR < 0.26
positively correlated
negatively correlated
H
EB-TCIP 24 h
Enrichment score
NES = 1.29
P-val < 0.03
FDR < 0.04
positively correlated
negatively correlated

**Figure SI-4: Global transcriptional changes induced by EB-TCIP**. Volcano plots of $log_2$fold changes of gene expression of cells treated with **BI3812** vs DMSO at 8 h (A) and 24 h (B), **NEG-1** vs DMSO at 8 h (C) and 24 h (D), and **EB-TCIP** vs **BI3812** (E) or **NEG-1** (F) at 24 h. **EB-TCIP** shows more dynamic changes in gene expression than either compound at both time points. At the 24 h time point *CISH* is not differentially expressed as seen in our time course. *SOCS2* was not differentially expressed in **EB-TCIP** vs **BI3812** during these experiments. Volcano plots are averages of three biological replicates for each treatment. GSEA shows a strong positive correlation between **EB-TCIP** induced gene expression and a published *BCL6* gene signature[1] at both 8 (G) and 24 h (H). RNA-seq data is shown as the average of three independent replicates.

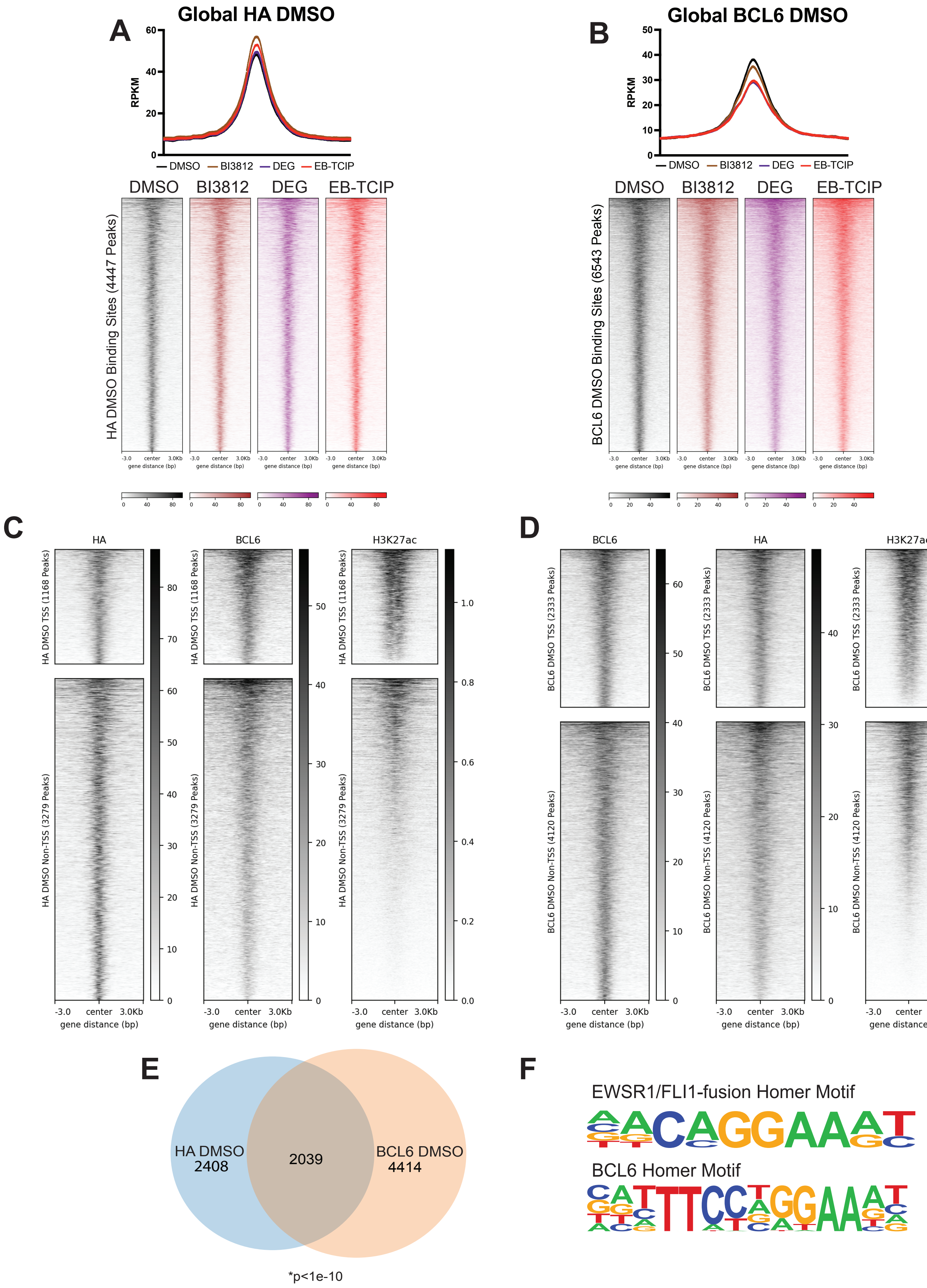
A
Global HA DMSO
RPKM
DMSO
BI3812
DEG
EB-TCIP
HA DMSO Binding Sites (4447 Peaks)
B
Global BCL6 DMSO
BCL6 DMSO Binding Sites (6543 Peaks)
C
HA
BCL6
H3K27ac
HA DMSO TSS (1168 Peaks)
HA DMSO Non-TSS (3279 Peaks)
D
BCL6 DMSO TSS (2333 Peaks)
BCL6 DMSO Non-TSS (4120 Peaks)
gene distance (bp)
E
HA DMSO
2408
2039
BCL6 DMSO
4414
*p<1e-10
F
EWSR1/FLI1-fusion Homer Motif
BCL6 Homer Motif

**Figure SI-5: FKBP-E/F and BCL6 share some binding sites on chromatin.** Global plot profiles (top) and volcano plots (bottom) for HA (A) and BCL6 (B) binding in DMSO (black), **BI3812** (brown), and **BI3802** (**DEG**, purple) treated samples. All compounds increase HA (FKBP-E/F) binding to chromatin to varying degrees. **BI3802 (DEG)** and **EB-TCIP** decrease BCL6 binding to chromatin, while **BI3812** has minimal effect. (C) HA DMSO ChIP-seq peaks projected into DMSO BCL6 ChIP-seq and H3K27ac ChIP-seq peaks from Lu, *et al.*[2]. (D) BCL6 DMSO ChIP-seq peaks projected into DMSO HA ChIP-seq and H3K27Ac ChIP-seq data from Lu, *et al.*[2]. (E) Venn diagram of overlapping ChIP-seq peaks in HA DMSO and BCL6 DMSO. These conditions share ~50% of their peaks. HA DMSO and BCL6 DMSO share more peaks than would be expected by chance based on a Fisher exact test. (F) BCL6 motif enrichment pattern and EWSR1/FLI1 fusion motif enrichment pattern. Both sequences contain the canonical "GGAA" EWS/FLI1 binding motif. All ChIP-seq is portrayed as the average of two independent replicates.

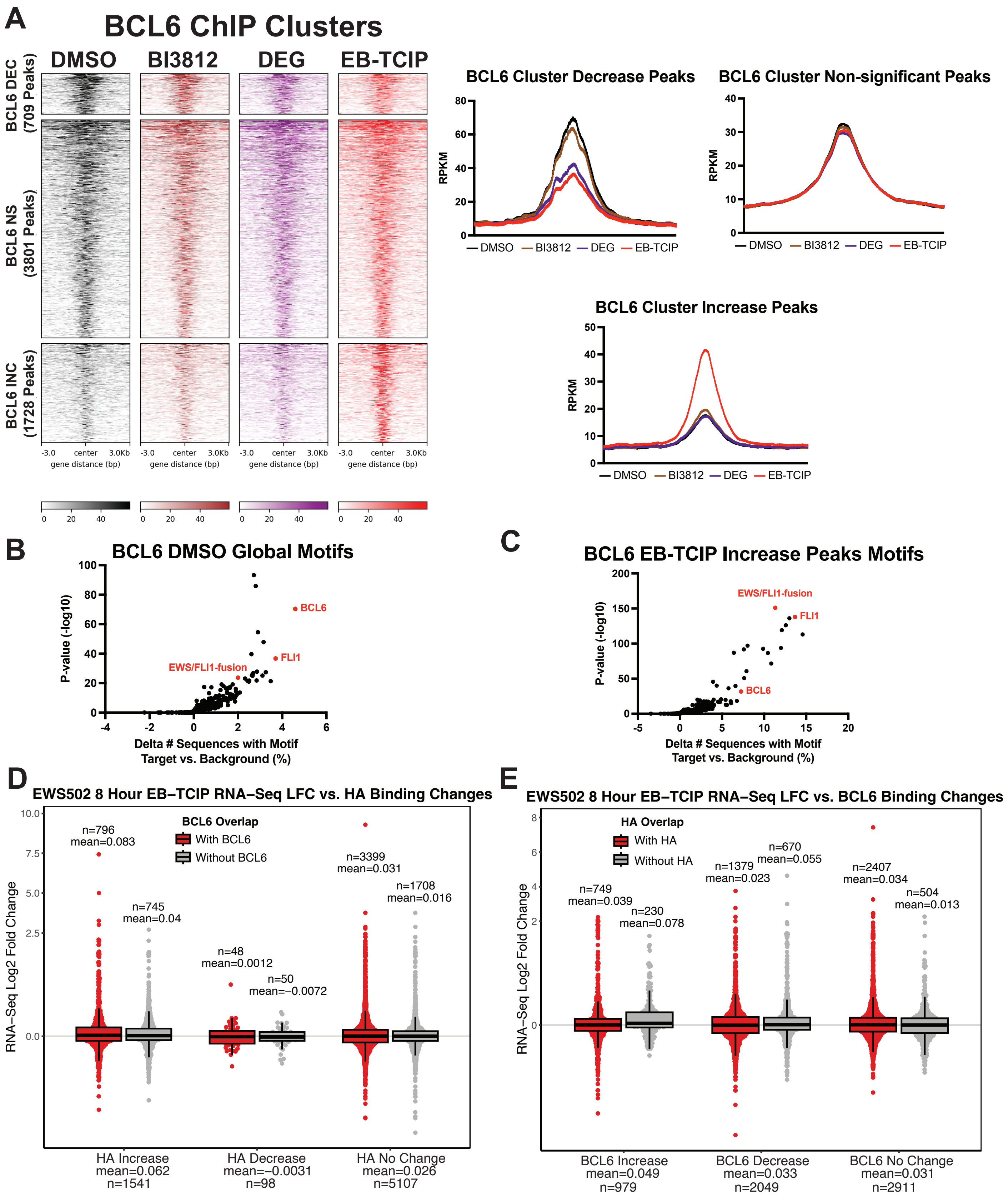


**Figure SI-6: EB-TCIP changes BCL6 binding on chromatin but does not have as much of an effect on global transcription as HA binding changes on chromatin.** (A) ChIP-seq tornado plots of

BCL6 binding signal for **EB-TCIP** versus DMSO peaks that are decreasing (DEC; 709), non-significantly changing (NS; 3801), and increasing (INC; 1728) in DMSO (black), **BI3812** (brown), and **BI3802** (**DEG**, purple) treated samples. Differential peaks between **EB-TCIP** and DMSO are shown for all compounds. **EB-TCIP** and **BI3802** show a loss of BCL6 binding at similar sites; however, only **EB-TCIP** induces an increase of BCL6 binding at a subset of peaks. Line plots for all compound treatments in each cluster are shown to the right. (B) Motif analysis of peaks in BCL6 DMSO. (C) Motif analysis of peaks in BCL6 **EB-TCIP** increased peaks; EWS/FLI1 peaks are enriched compared to BCL6 DMSO while the BCL6 motif decreases in rank. EWS502 8 h **EB−TCIP** vs. DMSO RNA−Seq $\log_2$fold change versus HA (FKBP−E/F) binding changes (D) or BCL6 binding changes (E) with **EB−TCIP** treatment. Stripchart plots show categories of EWS502 24 h **EB−TCIP** vs. DMSO HA (FKBP−E/F) or BCL6 ChIP−Seq differential peak analysis with strict AUC filtering on the x−axis with EWS502 8 h **EB−TCIP** vs. DMSO RNA−Seq DESeq2 log fold change on the non−linear y−axis (LFC>1 are compressed by a factor of 5). In (D) HA peak changes are split into subsets based upon whether they overlap with BCL6 whereas in (E) BCL6 peak changes are split into subsets based upon whether they overlap with HA. Peaks are mapped to nearest genes using Homer. Each point in the plot represents the $\log_2$fold change for a gene in EWS502 8 h **EB−TCIP** vs. DMSO DESeq2. The box plot shows median with first and third quartiles and the whiskers show 1.5 times the inter−quartile range. For p-value statistics comparing each subset for HA binding changes see Supporting Information Table 2. For p-value statistics comparing each subset for BCL6 binding changes see Supporting Information Table 3. All ChIP-seq is portrayed as the average of two independent replicates.

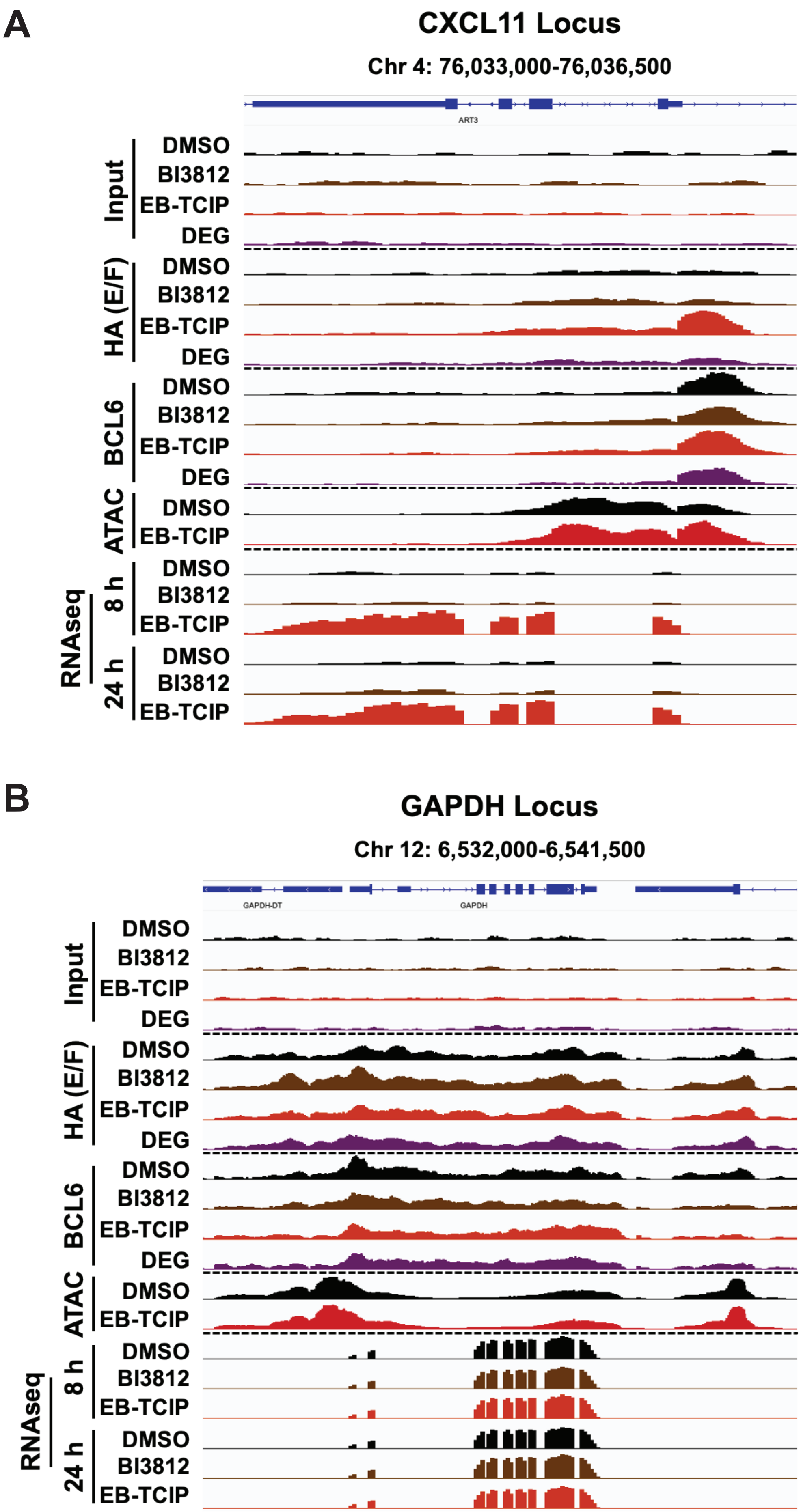
A
CXCL11 Locus
Chr 4: 76,033,000-76,036,500
ART3
Input
HA (E/F)
BCL6
ATAC
RNAseq
8 h
24 h
DMSO
BI3812
EB-TCIP
DEG
B
GAPDH Locus
Chr 12: 6,532,000-6,541,500
GAPDH-DT
GAPDH

**Figure SI-7: FKBP-E/F binding increases at BCL6 target loci, but not unrelated loci.** IGV visualization of Input, HA (FKBP-E/F), BCL6, ATAC-seq signal, and RNA-seq signal at the *CXCL11* (A) and *GAPDH* (B) loci with treatments DMSO (black), 1 μM **BI3812** (brown), 1 μM **BI3802** (**DEG,** purple), and 1 μM **EB-TCIP** (red). All ChIP-seq and ATAC-seq is portrayed as the average of two independent replicates. RNA-seq is portrayed as the average of three independent replicates.

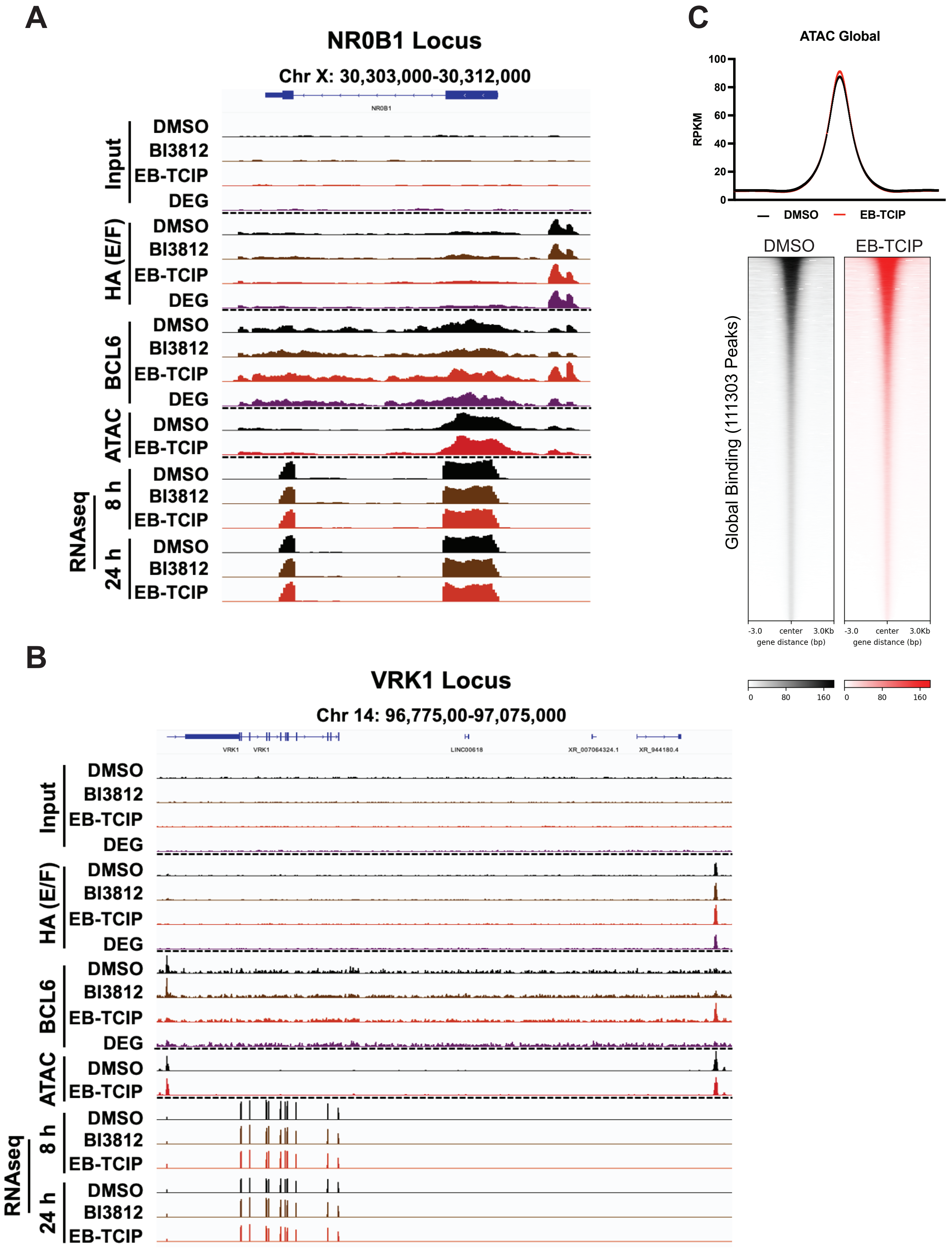
A
NR0B1 Locus
Chr X: 30,303,000-30,312,000
NR0B1
Input
HA (E/F)
BCL6
ATAC
RNAseq
8 h
24 h
DMSO
BI3812
EB-TCIP
DEG
B
VRK1 Locus
Chr 14: 96,775,00-97,075,000
VRK1
LINC00618
XR_007064324.1
XR_944180.4
C
ATAC Global
RPKM
100
80
60
40
20
0
DMSO
EB-TCIP
Global Binding (111303 Peaks)
-3.0
center
3.0Kb
gene distance (bp)
0
80
160

**Figure SI-8: EB-TCIP moves BCL6 to EWS/FLI1 sites with minimal effect on chromatin accessibility and expression.** IGV visualization of Input, HA (FKBP-E/F), BCL6, ATAC-seq signal, and RNA-seq signal at the *NR0B1* (A) and *VRK1* (B) with treatments DMSO (black), 1 μM **BI3812** (brown), 1 μM **BI3802** (**DEG**, purple), and 1 μM **EB-TCIP** (red). (C) Global ATAC-seq signal in DMSO (black) vs **EB-TCIP** (red) displayed as a summary plot (top) and tornado plot (bottom). **EB-TCIP** has minimal effects on global chromatin accessibility. All ChIP-seq and ATAC-seq is portrayed as the average of two independent replicates. RNA-seq is portrayed as the average of three independent replicates.

### Chemical Synthesis

#### Synthesis of EB-TCIP (BAK-04-212)

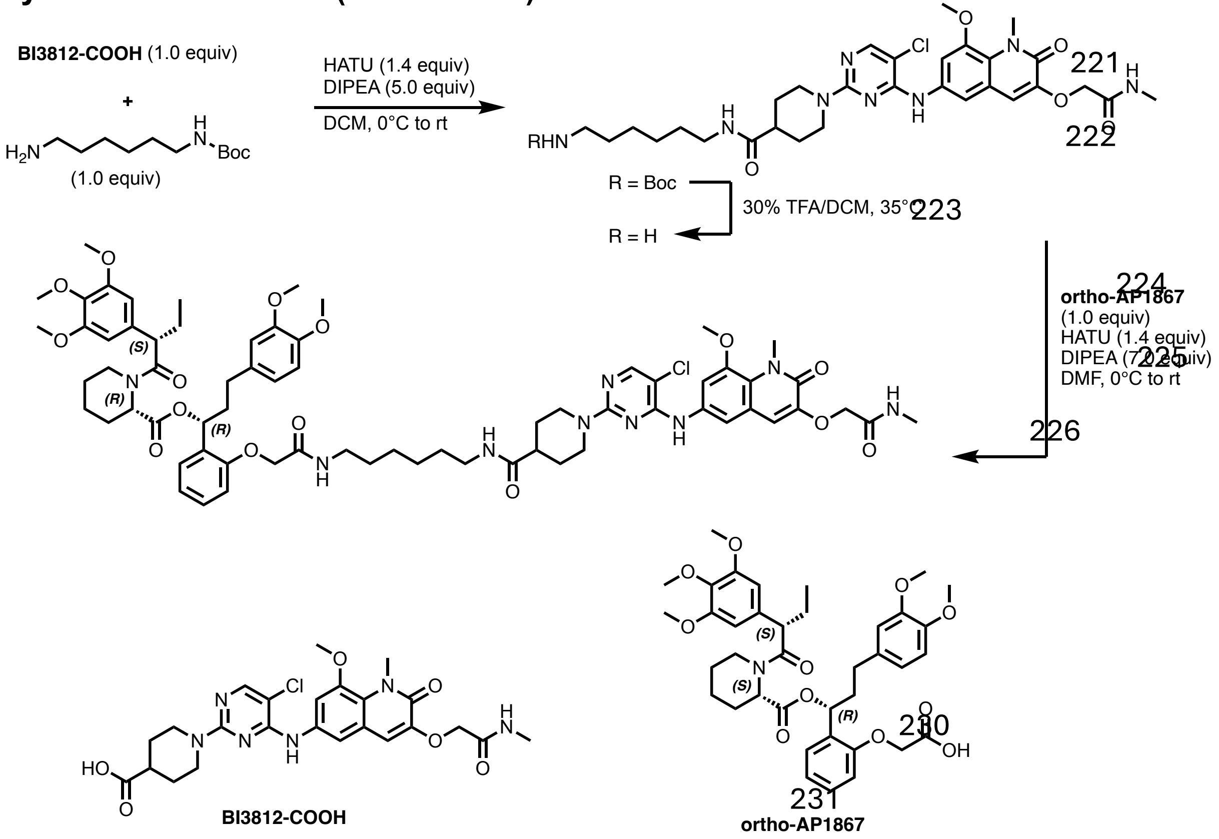


To a solution of **BI3812-COOH**[3] (30.0 mg, 0.057 mmol, 1.0 equiv), HATU (30.0 mg, 0.080 mmol, 1.4 equiv) and DIPEA (0.05 mL, 0.28 mmol, 5.0 equiv) in DCM (0.4 mL) at 0 °C was added *tert*-butyl (6-aminohexyl)carbamate (12.0 mg, 0.057 mmol, 1.0 equiv). The mixture was allowed to warm to warm to room temperature and, after stirring for 1 h, LC-MS analysis indicated full consumption of starting material. The mixture was then diluted with water and the aqueous layer was extracted with DCM (3x). The combined organic extracts were washed with brine, dried over $MgSO_4$, and concentrated under reduce pressure. The resulting crude material was purified via flash column chromatography on silica (0-10% MeOH in DCM) to afford the intermediate (9.8 mg, 24% yield). The intermediate was taken up in DCM (0.2 mL), TFA (0.06 mL) was added, and the orange solution was stirred at 35 °C for 1 h, at which point LC-MS analysis indicated clean removal of the Boc protecting group. The mixture was concentrated under reduced pressure and dried under high vacuum to remove excess TFA. Then, a solution of the residue in DMF (0.1 mL) was slowly added to a solution of **ortho-AP1867**[4] (9.4 mg, 0.014 mmol, 1.0 equiv), HATU (7.2 mg, 0.019 mmol, 1.4 equiv), and DIPEA (0.016 mL, 0.095 mmol,

7.0 equiv) in DMF (0.2 mL) at 0 °C. The mixture was allowed to warm to ambient temperature and after stirring for 1 h was directly purified by reverse-phase HPLC (40–100% MeCN in water, no acidic additive) to afford **EB-TCIP** as off-white solid upon lyophilization (8.1 mg, 46% yield).

**$^1$H NMR** (500 MHz, DMSO-d$_6$) Major Peaks: δ = 8.77 (s, 1H), 8.05 (s, 1H), 7.94 (d, *J* = 4.9 Hz, 1H), 7.73 (t, *J* = 5.5 Hz, 1H), 7.69 (t, *J* = 5.8 Hz, 1H), 7.54 (t, *J* = 1.9 Hz, 2H), 7.19 (m, 1H), 6.99 (s, 1H), 6.89 – 6.70 (m, 5H), 6.65 – 6.58 (m, 2H), 6.55 (s, 2H), 6.02 (dd, *J* = 8.4, 4.9 Hz, 1H), 5.38 – 5.29 (m, 1H), 4.63 – 4.45 (m, 6H), 4.04 (d, *J* = 13.2 Hz, 1H), 3.86 (assigned by HSQC, 7H), 3.71 (s, 3H), 3.69 (s, 3H), 3.56 (s, 6H), 3.54 (s, 3H), 3.15 – 3.02 (m, 2H), 2.98 (q, *J* = 6.4 Hz, 2H), 2.87 (t, *J* = 12.2 Hz, 2H), 2.64 (d, *J* = 4.7 Hz, 3H), 2.62 – 2.55 (m, 1H), 2.46 (assigned by HSQC, 1H), 2.42 – 2.31 (m, 2H), 2.14 (d, *J* = 14.4 Hz, 1H), 2.08 – 1.80 (m, 4H), 1.69 (d, *J* = 12.6 Hz, 2H), 1.63 – 1.44 (m, 7H), 1.32 (m, 4H), 1.28 – 1.09 (m, 4H), 0.86 – 0.75 (m, 3H).

**LC-MS**: m/z 653.70 [M+2]$^{2+}$.

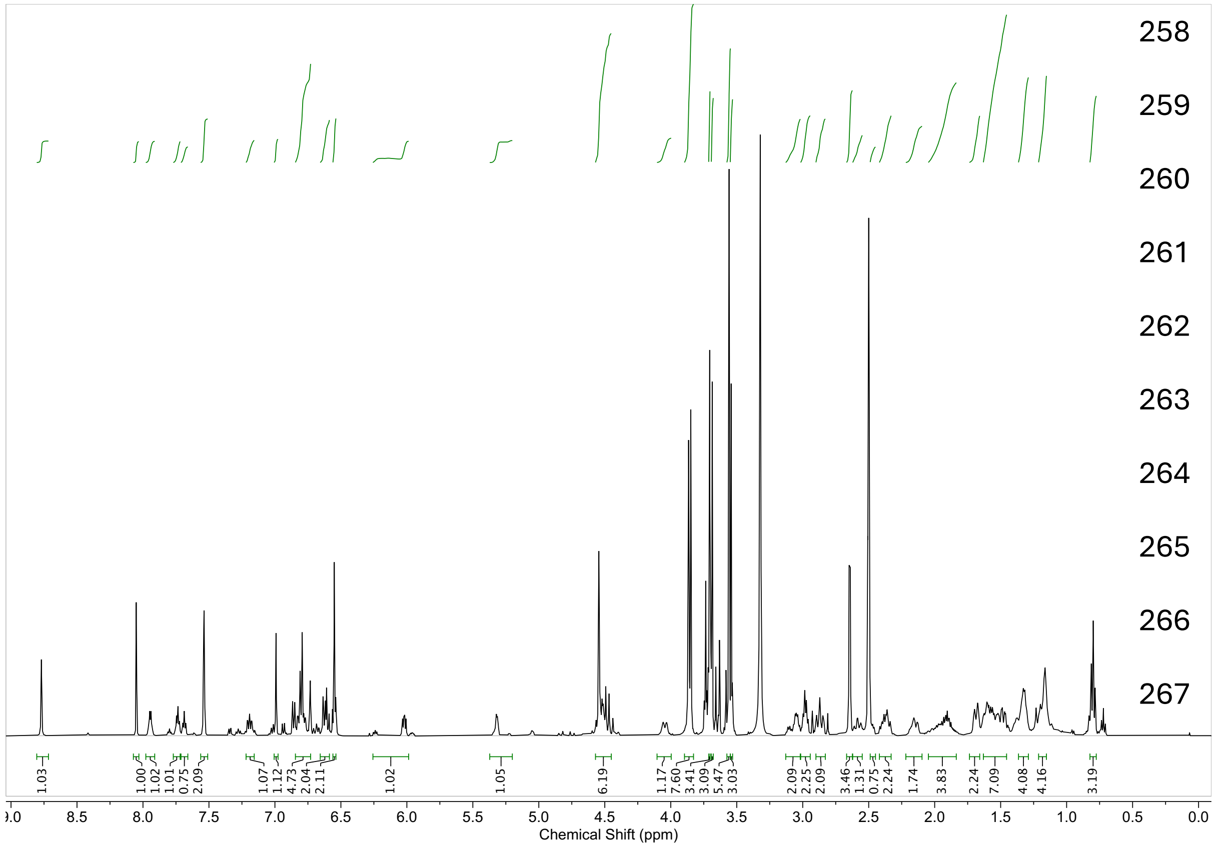

### Synthesis of NEG-1 (RPG-02-089)

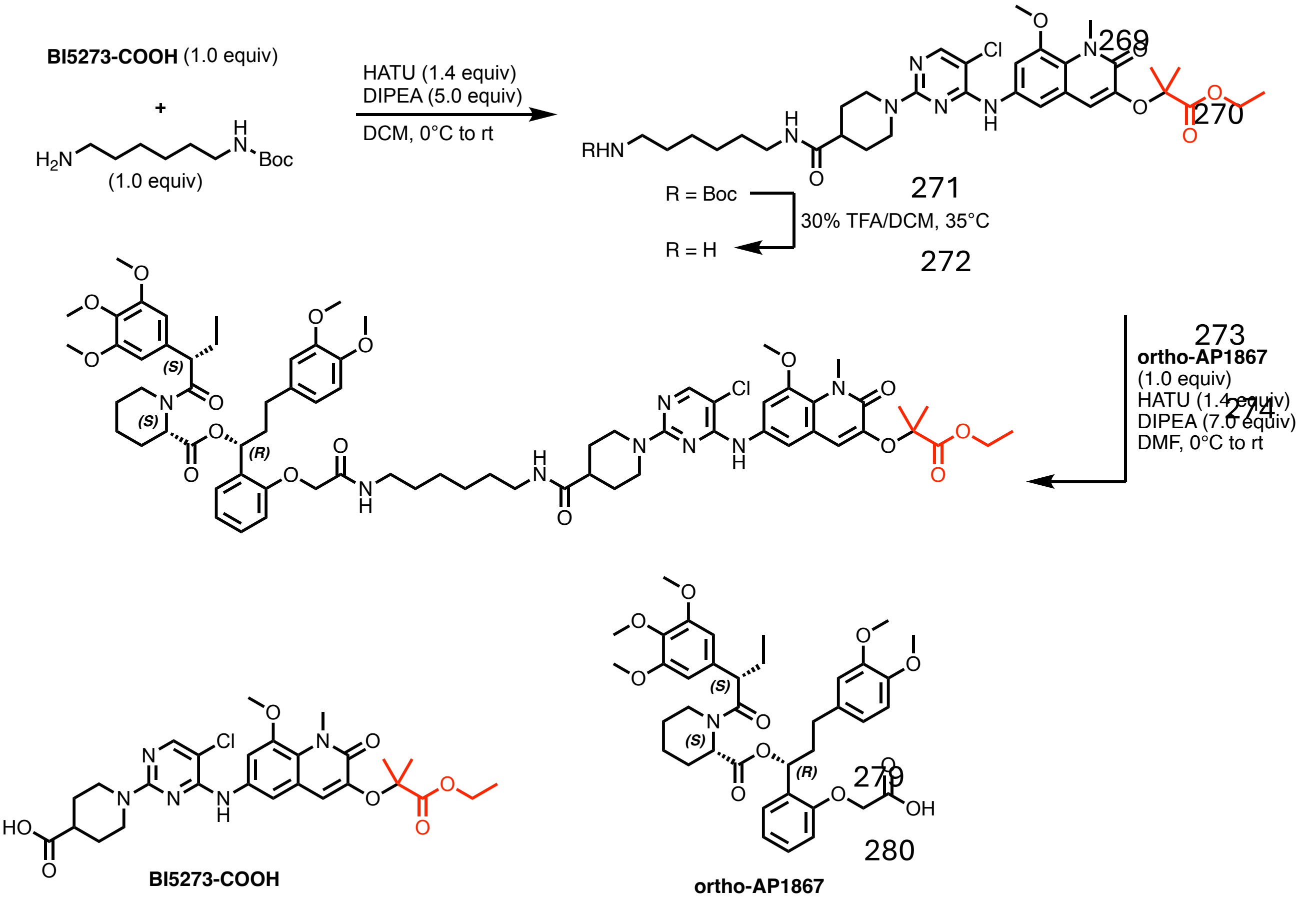


To a solution of **BI5723-COOH**[3] (20.0 mg, 0.035 mmol, 1.0 equiv), HATU (19.0 mg, 0.049 mmol, 1.4 equiv) and DIPEA (0.030 mL, 0.17 mmol, 5.0 equiv) in DCM (0.4 mL) at 0 °C was added *tert*-butyl (6-aminohexyl)carbamate (7.5 mg, 0.035 mmol, 1.0 equiv). The mixture was allowed to warm to warm to room temperature and, after stirring for 1 h, LC-MS analysis indicated full consumption of starting material. The mixture was then diluted with water and the aqueous layer was extracted with DCM (3x). The combined organic extracts were washed with brine, dried over $MgSO_4$, and concentrated under reduce pressure. The resulting crude material was purified via flash column chromatography on silica (0-10% MeOH in DCM) to afford the intermediate (21.3 mg, 79% yield). The intermediate was taken up in DCM (0.3 mL), TFA (0.09 mL) was added, and the orange solution was stirred at 35 °C for 1 h, at which point LC-MS analysis indicated clean removal of the Boc protecting group. The mixture was concentrated under reduced pressure and dried under high vacuum to remove excess TFA. Then, a solution of the residue in DMF (0.1 mL) was slowly added to a solution of **ortho-AP1867**[4] (19.0 mg, 0.028 mmol, 1.0 equiv), HATU (15.0 mg, 0.039 mmol, 1.4 equiv), and DIPEA (0.034 mL, 0.193 mmol,

7.0 equiv) in DMF (0.2 mL) at 0 °C. The mixture was allowed to warm to ambient temperature and after stirring for 1 h was directly purified by reverse-phase HPLC (40–100% MeCN in water, no acidic additive) to afford **NEG-1** as off-white solid upon lyophilization (6.8 mg, 18% yield).

**$^1$H NMR** (500 MHz, DMSO-$d_6$) Major Peaks: δ = 8.75 (s, 1H), 8.05 (s, 1H), 7.74 (t, J = 5.5 Hz, 1H), 7.68 (t, J = 5.8 Hz, 1H), 7.60 (d, J = 2.3 Hz, 1H), 7.52 (d, J = 2.2 Hz, 1H), 7.19 (ddd, J = 8.6, 6.5, 2.5 Hz, 1H), 6.86 (d, *J* = 8.3 Hz, 1H), 6.83 – 6.76 (m, 4H), 6.73 (d, J = 2.0 Hz, 1H), 6.69 – 6.57 (m, 2H), 6.55 (s, 2H), 6.02 (dd, J = 8.3, 4.9 Hz, 1H), 5.37 – 5.28 (m, 1H), 4.61 – 4.39 (m, 4H), 4.15 (q, J = 7.1 Hz, 2H), 4.04 (d, J = 13.1 Hz, 1H), 3.86 (assigned by HSQC, 4H), 3.81 (s, 3H), 3.70 (s, 3H), 3.69 (s, 3H), 3.56 (assigned by HSQC, 6H), 3.54 (assigned by HSQC, 3H), 3.15 – 3.02 (m, 2H), 2.98 (q, J = 6.4 Hz, 2H), 2.88 (t, J = 12.5 Hz, 2H), 2.65 – 2.55 (m, 1H), 2.48 – 2.43 (m, 1H), 2.44 – 2.31 (m, 2H), 2.14 (d, J = 13.5 Hz, 1H), 2.02 – 1.81 (m, 3H), 1.69 (d, J = 12.6 Hz, 2H), 1.55 (assigned by HSQC, 12H), 1.41 – 1.26 (m, 5H), 1.15 (m, 8H), 0.80 (t, J = 7.3 Hz, 3H).

**LC-MS**: m/z 675.25 $[M+2]^{2+}$.

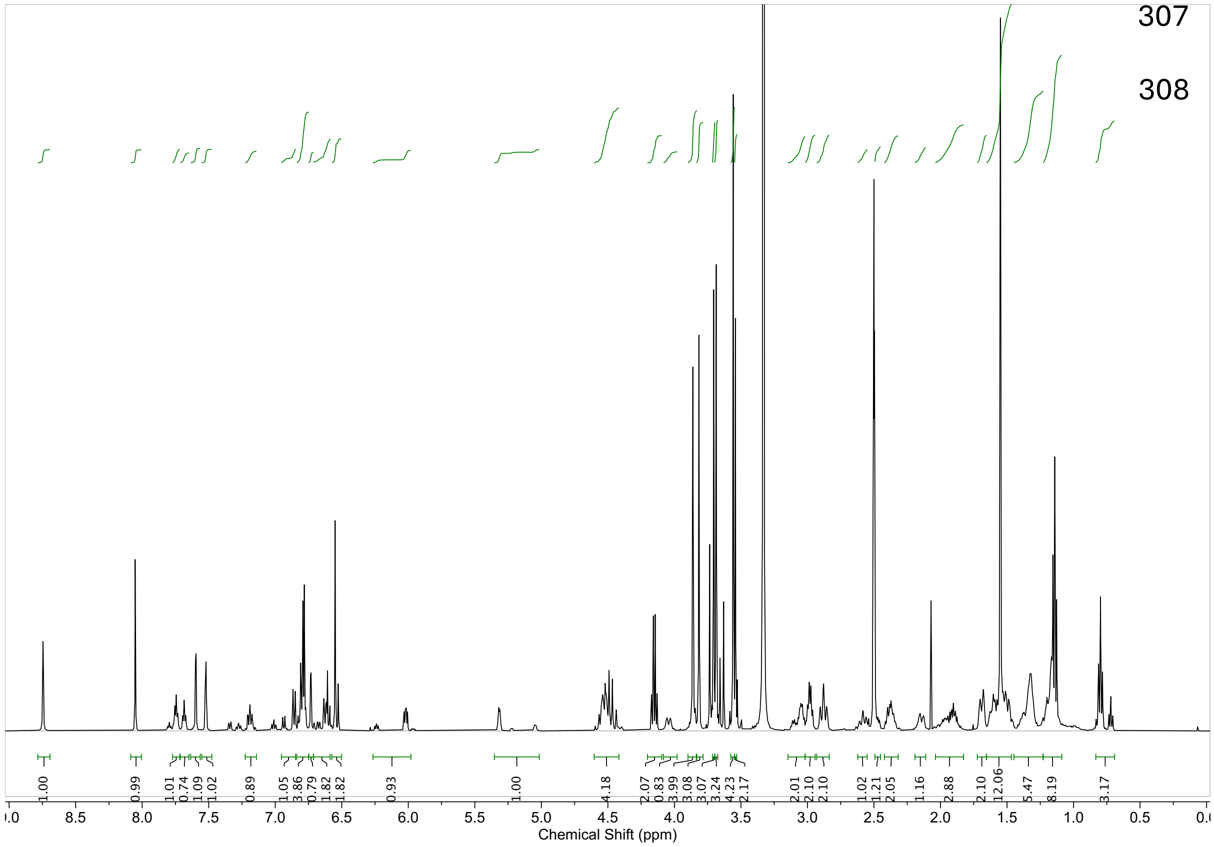

### Synthesis of NEG-2 (RPG-02-205)

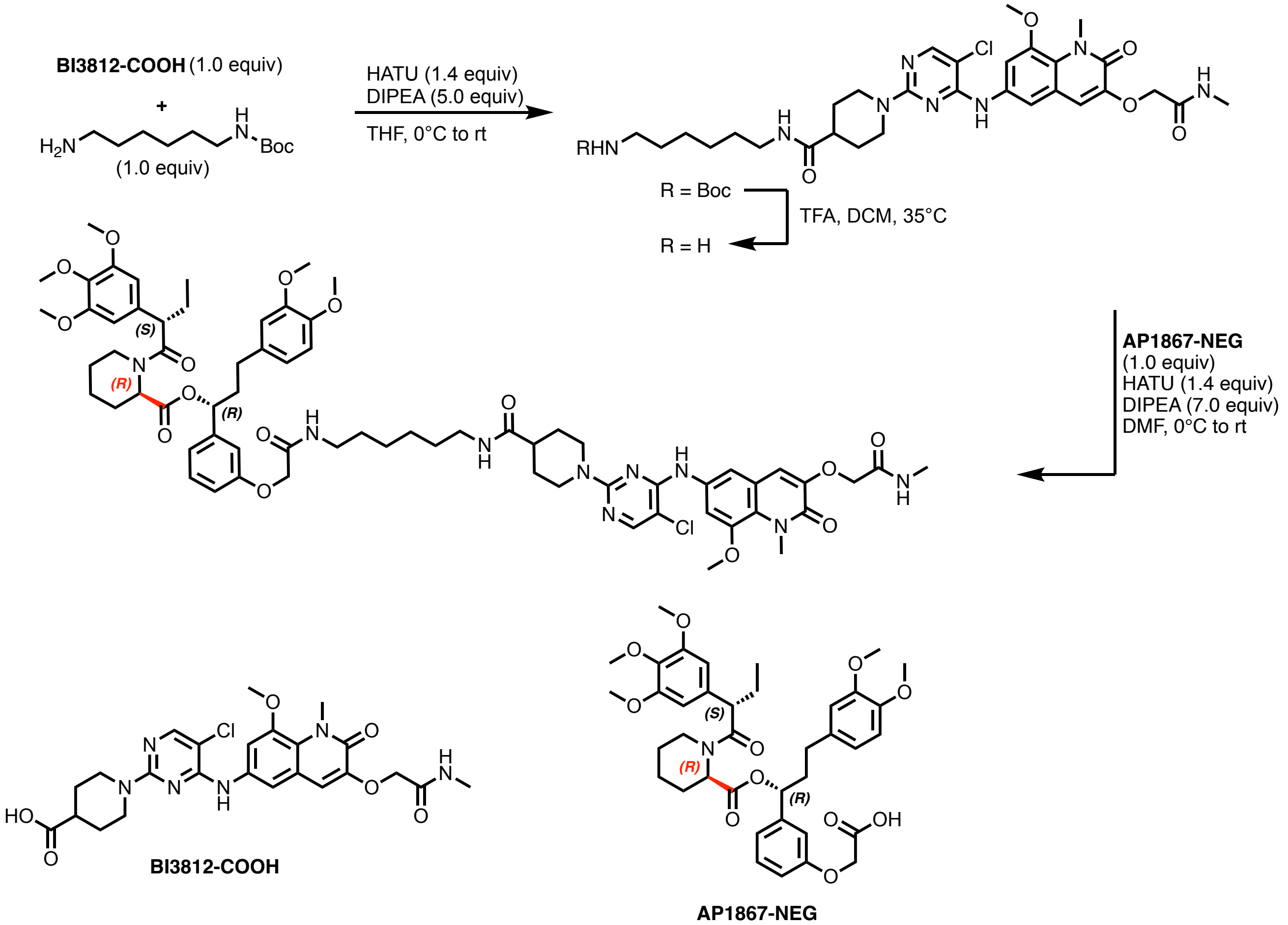


To a solution of **BI3812-COOH**[3] (32.0 mg, 0.06 mmol, 1.0 equiv), HATU (32.0 mg, 0.084 mmol, 1.4 equiv) and DIPEA (0.052 mL, 0.3 mmol, 5.0 equiv) in THF (0.4 mL) at 0 °C was added *tert*-butyl (6-aminohexyl)carbamate (13.0 mg, 0.057 mmol, 1.0 equiv). The mixture was allowed to warm to warm to room temperature and, after stirring for 1 h, LC-MS analysis indicated full consumption of starting material. The mixture was then diluted with water and the aqueous layer was extracted with DCM (3x). The combined organic extracts were washed with brine, dried over $MgSO_4$, and concentrated under reduced pressure. The resulting crude material was purified via flash column chromatography on silica (0-10% MeOH in DCM) to afford the intermediate (11.9 mg, 27% yield). The intermediate was taken up in DCM (0.2 mL), TFA (0.06 mL) was added, and the orange solution was stirred at 35 °C for 1 h, at which point LC-MS analysis indicated clean removal of the Boc protecting group. The mixture was concentrated under reduced pressure and dried under high vacuum to remove excess TFA. Then, a solution of the residue in DMF (0.25 mL) was slowly added to a solution of **AP1867-NEG**[5] (11.0 mg, 0.016 mmol, 1.0 equiv), HATU (8.5 mg, 0.022 mmol, 1.4 equiv), and DIPEA (0.019 mL, 0.11 mmol, 7.0

equiv) in DMF (0.25 mL) at 0 °C. The mixture was allowed to warm to ambient temperature and after stirring for 1 h was directly purified by reverse-phase HPLC (40–100% MeOH in water +0.035% TFA) to afford **NEG-2** as off-white solid upon lyophilization (6.3 mg, 30% yield).

**$^1$H NMR** (500 MHz, DMSO-$d_6$) Major Peaks: δ = 8.95 (s, 1H), 8.08 (s, 1H), 8.03 (dd, J = 11.5, 6.0 Hz, 1H), 7.95 (d, J = 4.9 Hz, 1H), 7.76 (t, J = 5.6 Hz, 1H), 7.52 (s, 2H), 7.27 (td, J = 8.1, 3.0 Hz, 1H), 7.00 (s, 1H), 6.93 – 6.81 (m, 4H), 6.76 (dd, J = 4.9, 2.1 Hz, 1H), 6.69 (m, 2H), 6.61 (s, 2H), 5.64 (dd, J = 8.3, 5.1 Hz, 1H), 5.37 – 5.30 (m, 1H), 4.55 (s, 2H), 4.48 (d, J = 13.4 Hz, 2H), 4.45 (s, 2H), 4.06 – 4.00 (m, 1H), 3.90 – 3.82 (assigned by HSQC, 7H), 3.78 – 3.72 (assigned by HSQC, 9H), 3.71 – 3.69 (assigned by HSQC, 3H), 3.63 (assigned by HSQC, 3H), 3.11 – 2.97 (m, 5H), 2.90 (t, J = 12.5 Hz, 2H), 2.64 (d, J = 4.7 Hz, 3H), 2.62 – 2.53 (m, 2H), 2.41 – 2.31 (m, 1H), 2.12 (d, J = 13.9 Hz, 2H), 2.06 – 1.89 (m, 2H), 1.70 (d, J = 12.6 Hz, 2H), 1.63 – 1.48 (m, 5H), 1.46 – 1.31 (m, 5H), 1.20 (m, 4H), 1.04 (d, J = 13.1 Hz, 1H), 0.89 – 0.76 (m, 4H).

**LC-MS**: m/z 653.40 $[M+2]^{2+}$.

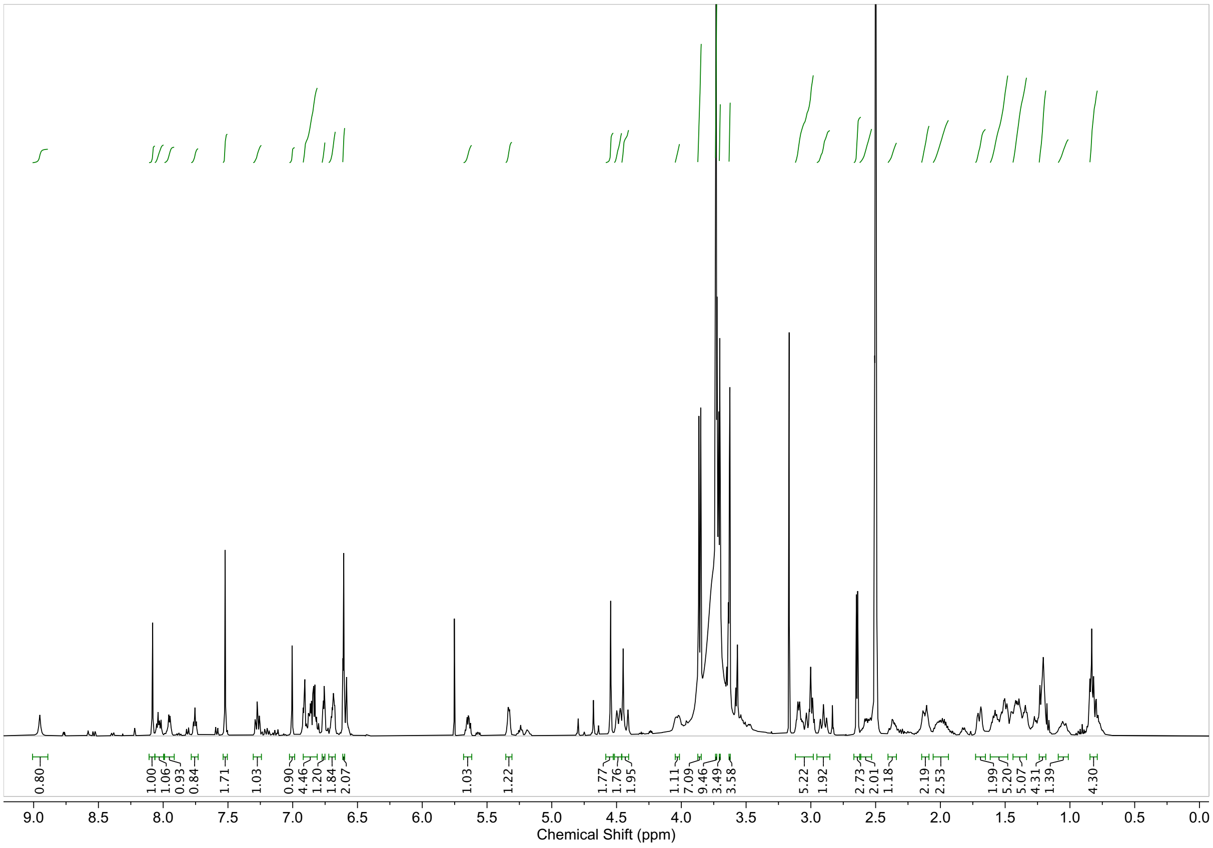

**Supporting Information Tables:**

| Cell Line | SOCS2 EC50 (nM) | Max SOCS2 Expression Ratio EB-TCIP:BI3812 | CISH EC50 (nM) | Max CISH Expression Ratio EB-TCIP:BI3812 |
|---|---|---|---|---|
| **EWS502** | 172 ± 47 | 3.3 ± 0.6 | 107 ± 44 | 2.1 ± 0.4 |
| **TC32** | 102 ± 12 | 2.4 ± 0.7 | 118 ± 65 | 2.5 ± 1.1 |

**SI-Table 1: EB-TCIP activity in FKBP-E/F expressing EWS502 and TC32.** The $EC_{50}$ of **EB-TCIP** is in the mid nanomolar range in both cell lines for both *SOCS2* and *CISH*. **EB-TCIP** induces 2-3 fold higher expression of these genes than **BI3812** in both cell lines.

| | HA.Inc.w.BCL6 | HA.Inc.wo.BCL6 | HA.Dec.w.BCL6 | HA.Dec.wo.BCL6 | HA.No.Change.w.BCL6 |
|---|---|---|---|---|---|
| *HA Inc wo BCL6* | 0.053 | NA | NA | NA | NA |
| *HA Dec w BCL6* | 0.015 | 0.18 | NA | NA | NA |
| *HA Dec wo BCL6* | 0.0018 | 0.053 | 0.77 | NA | NA |
| *HA N.C. w BCL6* | 0.01 | 0.45 | 0.27 | 0.085 | NA |
| *HA N.C. wo BCL6* | 0.0016 | 0.065 | 0.56 | 0.27 | 0.11 |

**SI-Table 2: Significance table for Figure SI-6D:** p−values from a paired t−test with a Benjamini−Hochberg correction for the differences between RNA−seq mean LFC for the various subsets for **EB-TCIP** vs DMSO RNA-seq at 8 h.

| | BCL6.Inc.w.HA | BCL6.Inc.wo.HA | BCL6.Dec.w.HA | BCL6.Dec.wo.HA | BCL6.No.Change.w.HA |
|---|---|---|---|---|---|
| *BCL6 Inc wo HA* | 0.1 | NA | NA | NA | NA |
| *BCL6 Dec w HA* | 0.31 | 0.0087 | NA | NA | NA |
| *BCL6 Dec wo HA* | 0.42 | 0.31 | 0.092 | NA | NA |
| *BCL6 N.C. w HA* | 0.69 | 0.037 | 0.31 | 0.24 | NA |
| *BCL6 N.C. wo HA* | 0.16 | 0.0061 | 0.45 | 0.04 | 0.15 |

**SI-Table 3: Significance table for Figure SI-6E:** p−values from a paired t−test with a Benjamini−Hochberg correction for the differences between RNA−seq mean LFC for the various subsets for **EB-TCIP** vs DMSO RNA-seq at 8 h.

**Supporting Information References:**


1 Ci, W. *et al.* The BCL6 transcriptional program features repression of multiple oncogenes in primary B cells and is deregulated in DLBCL. *Blood* **113**, 5536-5548, doi:10.1182/blood-2008-12-193037 (2009).

2 Lu, D. Y. *et al.* The ETS transcription factor ETV6 constrains the transcriptional activity of EWS-FLI to promote Ewing sarcoma. *Nat Cell Biol* **25**, 285-297, doi:10.1038/s41556-022-01059-8 (2023).

3 Gourisankar, S. *et al.* Rewiring cancer drivers to activate apoptosis. *Nature* **620**, 417-425, doi:10.1038/s41586-023-06348-2 (2023).

4 Nabet, B. *et al.* The dTAG system for immediate and target-specific protein degradation. *Nat Chem Biol* **14**, 431-441, doi:10.1038/s41589-018-0021-8 (2018).

5 Hu, Z. *et al.* EGFR targeting PhosTACs as a dual inhibitory approach reveals differential downstream signaling. *Sci Adv* **10**, eadj7251, doi:10.1126/sciadv.adj7251 (2024).
